# Evaluating Methods of Correcting for Multiple Comparisons Implemented in SPM12 in Social Neuroscience fMRI Studies: An Example from Moral Psychology

**DOI:** 10.1101/129734

**Authors:** Hyemin Han, Andrea L. Glenn

## Abstract

In fMRI research, the goal of correcting for multiple comparisons is to identify areas of activity that reflect true effects, and thus would be expected to replicate in future studies. Finding an appropriate balance between trying to minimize false positives (Type I error) while not being too stringent and omitting true effects (Type II error) can be challenging. Furthermore, the advantages and disadvantages of these types of errors may differ for different areas of study. In many areas of social neuroscience that involve complex processes and considerable individual differences, such as the study of moral judgment, effects are typically smaller and statistical power weaker, leading to the suggestion that less stringent corrections that allow for more sensitivity may be beneficial, but also result in more false positives. Using moral judgment fMRI data, we evaluated four commonly used methods for multiple comparison correction implemented in SPM12 by examining which method produced the most precise overlap with results from a meta-analysis of relevant studies and with results from nonparametric permutation analyses. We found that voxel-wise thresholding with family-wise error correction based on Random Field Theory provides a more precise overlap (i.e., without omitting too few regions or encompassing too many additional regions) than either clusterwise thresholding, Bonferroni correction, or false discovery rate correction methods.

---

Evaluating Methods of Correcting for Multiple Comparisons Implemented in SPM12 in Social Neuroscience fMRI Studies: An Example from Moral Psychology

Correcting for multiple comparisons has been one of the most significant challenges in the statistical analysis of fMRI data (Bennett, Miller, & Wolford, 2009). Because more than one hundred thousand voxels are compared simultaneously during analysis, the chances of Type I error are very high in the absence of any correction (Genovese, Lazar, & Nichols, 2002). In order to address this issue, researchers have developed various correction methods. For instance, Bonferroni’s correction method, one of the traditional methods for multiple comparison correction, divides the nominal significance level (e.g., *p* < .05) by the number of tests being performed (Bland & Altman, 1995). Although Bonferroni correction produces good control of Type I error, it has the disadvantage of removing both false and true positives when applied to whole brain analyses. To address this issue, many researchers use a family-wise error (FWE) correction method based on Random Field Theory (RFT) (Nichols, 2012). Unlike the traditional Bonferroni method, which only accounts for the total number of comparisons, this method assumes that the error fields can be a lattice approximation to an underlying random field usually with a Gaussian distribution (Brett, Penny, & Kiebel, 2004; Eklund, Nichols, & Knutsson, 2016). Moreover, the false discovery rate (FDR) correction method was developed. This method is thought to be more sensitive and less likely to produce Type II error than FWE correction methods. Unlike the aforementioned methods that control for the possibility of any false positives, this method focuses on the expected proportion of false positives only among survived entities (Genovese et al., 2002; Nichols, 2013). In terms of the implementation of the FDR correction method, neuroimaging has relied on the standard FDR procedure, the linear step-up procedure, or so-called Benjamini and Hochberg procedure (Benjamini & Hochberg, 1995; Benjamini, Krieger, & Yekutieli, 2006), although more sophisticated procedures, such as adaptive linear step-up procedures, have been developed (Benjamini et al., 2006).

These correction methods can be performed at different levels of inference (i.e., voxel-wise and clusterwise inference; Flandin & Novak (2013) using fMRI analysis software (e.g., SPM) and customized MATLAB codes. In the case of voxel-wise inference, each individual voxel is treated as a unit for analysis, and any voxel exceeding a threshold after applying one of the aforementioned correction methods is considered statistically significant in the whole brain or specified regions of interest (Nichols, 2012). In the case of clusterwise inference, statistically significant clusters showing activation are detected based on the number of contiguous voxels; this type of inference does not control the estimated false positive probability of each individual voxel in each region, but controls such a probability of the region as a whole (Woo, Krishnan, & Wager, 2014). This clusterwise inference has been one of the most popular methods used for multiple comparison correction because it is considered to be more sensitive than voxel-wise inference (Woo et al., 2014).

Although the aforementioned correction methods, particularly RFT FWE correction and FDR correction, have been implemented in widely used fMRI analysis software (e.g., SPM) with parametric assumptions, a nonparametric analysis tool, Statistical non-Parametric Mapping (SnPM) (Nichols & Holmes, 2002), uses permutations in order to correct for multiple comparisons without several assumptions required for parametric analysis, such as normally distributed data and mean parameterization. Instead, SnPM requires several minimal assumptions pertaining to the empirical null hypothesis (Nichols, 2012). For instance, in the case of a two-sample t-test, the subjects are assumed to be exchangeable under the null hypothesis, which might be violated if the subjects are related; in the case of a one-sample t-test, sign flipping, which is based on an assumption that the errors have a symmetric distribution, is used (Winkler, Ridgway, Webster, Smith, & Nichols, 2014). The application of SnPM is considered less stringent than the Bonferroni and RFT-FWE correction methods applied by software supporting parametric analysis when it is applied using voxel-wise inference (Eklund et al., 2016; Nichols & Hayasaka, 2003); however, clusterwise inference with SnPM is considered to be more stringent (Eklund et al., 2016). Furthermore, the randomise function in FSL and the corresponding function in the BROCCOLI software (Eklund, Dufort, Villani, & LaConte, 2014), which are based on the same statistical principles used for SnPM, have been utilized to evaluate false-positive rates and sensitivity of traditional correction methods (Eklund et al., 2016).

Although clusterwise inference has become a popular method for multiple comparison correction, a recent study evaluating different correction methods has raised concerns that the RFT-applied FWE correction method for clusterwise inference implemented in widely-used fMRI analysis software, such as SPM and FSL, inflates false-positive rates and produces erroneous outcomes (Eklund et al., 2016). RFT clusterwise inference relies on two strong assumptions that might cause such erroneous outcomes. It assumes that “the spatial smoothness of the fMRI signal is constant over the brain” (Eklund et al., 2016, p. 7902), and that there is a specific shape in the spatial autocorrelation function (Eklund et al., 2016). Using resting-state fMRI and cognitive experimental fMRI data analyzed with putative task designs, Eklund et al. (2016) report that rather than a false-positive rate of 5%, the most common software packages (SPM, FSL, AFNI) resulted in false-positive rates of up to 70% when RFT clusterwise inference was performed. However, the RFT-applied voxel-wise inference produced conservative outcomes with a false-positive rate of 5% or less (Eklund et al, 2016). Furthermore, there have been concerns regarding the FDR method as well. Although the FDR correction method reduces the probability of Type II error with enhanced sensitivity, this method is perhaps more likely to produce Type I error compared to other more conservative correction methods (Bennett, Wolford, & Miller, 2009).

The problem of appropriately correcting for multiple comparisons may be especially concerning in the field of social neuroscience in which balancing the risk of Type I and Type II error is sometimes an especially difficult task. Social and affective fMRI experiments often, though not always, produce smaller effect sizes and have weaker statistical power compared to, for example, sensory-motor experiments; there are several reasons for this (Lieberman & Cunningham, 2009). In many areas of social neuroscience, the psychological processes measured by these experiments are often poorly defined and are not directly observable (Poldrack, 2011). For example, in the field of moral psychology, the process of moral judgment – determining whether something is right or wrong –involves a number of different lower order processes, and is a task that can be accomplished in different ways by different individuals (Blasi & Hoeffel, 1974; Narvaez, Getz, Rest, & Thoma, 1999). Furthermore, the mental states of the individual are often not as certain. We cannot, for example, know with certainty that a person is “experiencing empathy” at a particular moment in time, nor expect the neural correlates of such a reported experience to be the same for all individuals. In contrast, with sensory and motor phenomena, there is a closer mapping between experimental inputs (e.g., a visual cue) and behavioral outputs (e.g., the person taps his fingers) and much less variability from trial-to-trial or person-to-person (Lieberman & Cunningham, 2009).

Although there are some examples of larger effects and more precise study designs within social neuroscience, such as the response of the fusiform face area to images of faces, many social neuroscience studies involve complex paradigms that may allow for multiple mental processes to occur, and in which the timing of processing is less precise. Thus, it has been argued that attempts to diminish Type I errors may be problematic in many social neuroscience studies, as they increase the probability of missing true effects (Lieberman & Cunningham, 2009). As a result, more lenient correction methods have become common in the field, in order to improve sensitivity. However, given the recent report demonstrating the possibility of inflated false positive rates possibly produced by the application of methods with better sensitivity (e.g., clusterwise inference), researchers should be cautious when applying such methods for correction in social neuroscientific studies.

Because of the unique goals and challenges of social neuroscience research, it is important to evaluate the various methods for multiple correction reviewed above in the context of a social neuroscientific experiment. Previous studies have evaluated different correction methods by using data from various clinical, visual, cognitive, and behavioral experiments (Eklund et al., 2016; Nichols & Hayasaka, 2003; Nichols & Holmes, 2002) and using simulation data (Nichols & Hayasaka, 2003), but not using data collected in a social neuroscience experiment. Because of the issues raised above regarding small effects and reduced statistical power, one of the challenges with many types of social neuroscience research is that it is more difficult to determine which activations are indeed false positives. However, in the context of the more complex and noisier data of some social neuroscience experiments, one way to generate an activation map that may more closely reflect the “true” pattern of activity for a particular process in order to evaluate these methods – is to use meta-analysis. Meta-analysis has been suggested as a feasible method to enhance the statistical power of fMRI analysis and better examine the overall brain activity pattern associated with a certain functionality of interest by analyzing a large amount of data while addressing the issue of reverse inference (Eklund et al., 2016; Lieberman & Cunningham, 2009; Poldrack, 2011).

Thus, in the present study we compare the quality of various correction methods in the analysis of data collected in one area of social neuroscience that is faced with the challenges mentioned above – an fMRI study of moral judgment. We used datasets and results from a quantitative meta-analysis of previously published fMRI studies on moral cognition and emotion (Han, 2017) in order to identify which correction method results in activations that most closely resemble results from the meta-analysis. Furthermore, as recommended by Nichols & Holmes (2002), we also compare results from each correction method with results produced by SnPM analyses, which does not require any parametric assumptions. We test four thresholding methods which have been widely used in the field – FDR, thresholding with RFT clusterwise inference, RFT-based FWE voxel-wise thresholding implemented in SPM, and Bonferroni correction.

## Methods and Materials

### Subjects and Materials

In the present study we reanalyzed previously collected fMRI data (Han, 2016; Han, Chen, Jeong, & Glover, 2016; Han, Glover, & Jeong, 2014). This fMRI data was obtained from 16 participants (8 male) who attended a Northern Californian university. They were undergraduate and graduate students who ranged in age from 21 to 34 years (*M* = 28.59, *SD* = 3.18). Functional brain images were acquired while they were making judgments about 60 socio-moral dilemmas that were previously developed for fMRI experiments (Greene, Nystrom, Engell, Darley, & Cohen, 2004; Greene, Sommerville, Nystrom, Darley, & Cohen, 2001). The dilemma set consisted of three different types of dilemmas: moral-personal (22 dilemmas), moral-impersonal (18 dilemmas), and non-moral dilemmas (20 dilemmas). Moral-personal dilemmas tend to provoke negative intuitive emotional responses among participants and often involve salient harm to human lives. Moral-impersonal dilemmas involved moral content but did not intend to provoke strong gut-level responses (e.g., should you return a lost wallet). Non-moral dilemmas included simple value-neutral mathematical problem sets. Participants were asked to choose one of two options to address each presented dilemma. Functional brain images were acquired using a spiral in-and-out sequence (TR = 2000ms, TE = 30ms, flip angle = 90) (Glover & Law, 2001). For the functional images, a total of 31 oblique axial slices were scanned parallel to the AC-PC with 4-mm slice thickness, 1-mm inter-slice skip. The resolution was 3.75 x 3.75 mm (FOV = 240mm, 64 x 64 matrix). Similar to Greene et al.’s (2001, 2004) experiments, we modeled brain activity four scans before, one during and three scans after the moment of response.

### Procedures

#### Reanalysis of fMRI data

We reanalyzed the brain images using four different approaches for multiple comparison correction. These approaches were FDR correction (Genovese et al., 2002), clusterwise inference-applied thresholding (Flandin & Novak, 2013), voxel-wise thresholding with FWE correction based on the RFT implemented in SPM12 (Flandin & Novak, 2013; Nichols, 2013), and FWE correction using Bonferroni’s method (Nichols & Holmes, 2002). Analyses were performed by using SPM12. The FDR inference was performed in MATLAB software and was based on example code developed by Nichols (2013). Also, a customized MATLAB code was composed to implement voxel-wise thresholding instead of peak-wise analysis that is the default in SPM. For the first-level estimation, a separate general linear model (GLM) was set for each participant that examined neural activity during each of three conditions. Each regressor was convolved with a standard hemodynamic response function (HRF). For comparisons between conditions, second-level estimation was conducted. The performed comparisons were conducted for these pairs: all moral (moral-personal + moral-impersonal) versus non-moral, moral-personal versus non-moral, moral-impersonal versus non-moral, moral-personal versus moral-impersonal, and moral-impersonal versus moral-personal. We applied four different thresholds for comparison. First, a voxel-wise threshold of *p* < .05 was used after applying one of the three multiple comparison correction methods (i.e., FDR, RFT FWE, Bonferroni’s FWE). Second, we also applied an uncorrected voxel-wise threshold of *p* < .001 with a clusterwise threshold of *p* < .05 after FWE correction, which was provided by SPM12. For cross-check, we also conducted the reanalysis with SnPM. Similarly, a voxel-wise threshold of *p* < .05 after applying FWE correction was used and 5,000 permutations were performed.

#### Meta-analysis of previous fMRI experiment for the basis for evaluation

We evaluated the four correction methods by comparing findings from our analyses of fMRI data to those from a meta-analysis of fMRI studies on moral cognition and emotion (Han, 2017). GingerALE software (version 2.3.6), which implements the activation likelihood estimation (ALE) method (Eickhoff et al., 2009; Eickhoff, Bzdok, Laird, Kurth, & Fox, 2012; Laird, Lancaster, & Fox, 2005), was employed in the meta-analysis. The meta-analysis examined a previously collected set of activation foci that were found by previous neuroimaging studies that compared neural correlates between moral and non-moral task conditions (for details see Han, 2017). This dataset included 45 experiments with 959 participants and 463 activation foci reported by 43 articles (see Table S1 for the list of included articles). In particular, comparisons between overall moral versus non-moral task conditions and moral versus non-moral judgment were performed. For the former comparison the whole dataset was meta-analyzed; for the comparison between moral versus non-moral judgment, we meta-analyzed a subset of the whole dataset (18 experiments with 373 participants and 142 activation foci reported in 17 articles) that included previous studies focusing on moral judgment among various moral functions. For both meta-analyses, we used FDR of. 01 as a cluster-forming threshold and. 05 for clusterwise inference as suggested (Fox et al., 2013). The calculated ALE map was compared with results produced by the aforementioned four correction methods for quality evaluation.

#### Overlap index calculation and quality evaluation

The present study aimed to quantitatively examine the overlap between survived activation foci after the application of each correction method and those found by the meta-analysis, foci in the ALE map created by GingerALE, and SnPM. We may simply represent the degree of overlap with the ratio of the number of overlapped voxels to the total number of voxels of a reference area. In case of the present study, two different types of ratios can be calculated: the ratio of the number of overlapped voxels (|*V*_*Ovl*_|) to the number of survived activation foci after applying correction method (|*V*_*Cor*_|; |*V*_*FDR*_| in case of FDR correction, |*V*_*CLU*_| in case of clusterwise inference thresholding, |*V*_*RFT*_| in case of RFT-applied FWE voxel-wise thresholding, |*V*_*Bon*_| in case of Bonferroni’s method-applied FWE correction) and that of the number of overlapped voxels to the number of activation foci found by meta-analysis (|*V*_*Meta*_|) and SnPM (|*V*_*SnPM*_|). However, these ratios would not provide us with enough information regarding the overall fit; instead, it may demonstrate a biased result. For instance, if survived voxels are completely contained by activation foci found by the meta-analysis, |*V*_*Ovl*_|*/*|*V*_*Cor*_| becomes 1.00, but |*V*_*Ovl*_|/|*V*_*Meta*_| can be smaller than 1.00. On the other hand, if resultant voxels from the meta-analysis are subsets of survived voxels, |*V*_*Ovl*_|*/*|*V*_*Meta*_| is 1.00 while |*V*_*Ovl*_|/|*V*_*Cor*_| can become smaller than 1.00. In these situations, we cannot make an accurate decision about which case shows a greater overlap solely based on the two independent ratios.

Instead, we may consider employing a unified overlap index, which takes into account both ratios simultaneously. The harmonic mean, instead of the arithmetic mean, would be a feasible and reliable way to calculate the overall overlap index with two different ratios, given its definition (Marchiori & Latora, 2000). We can then calculate the overall overlap index (*I*_*Ovl*_) as follows:

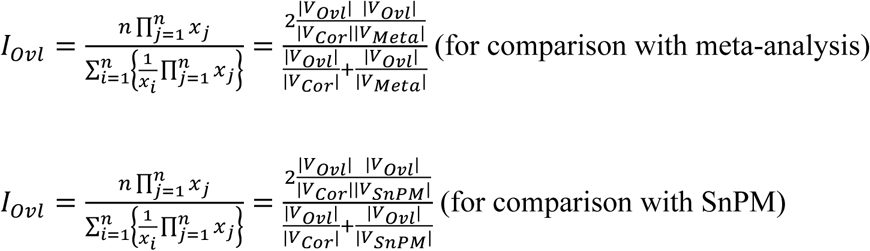

Five hypothetical cases are introduced as examples (see Figure 1). Red areas represent survived voxels after the application of correction method, yellow areas represent activation foci found by meta-analysis, and white areas represent overlapped voxels. Table 1 demonstrates two different ratios calculated for each case.

**Figure 1.**
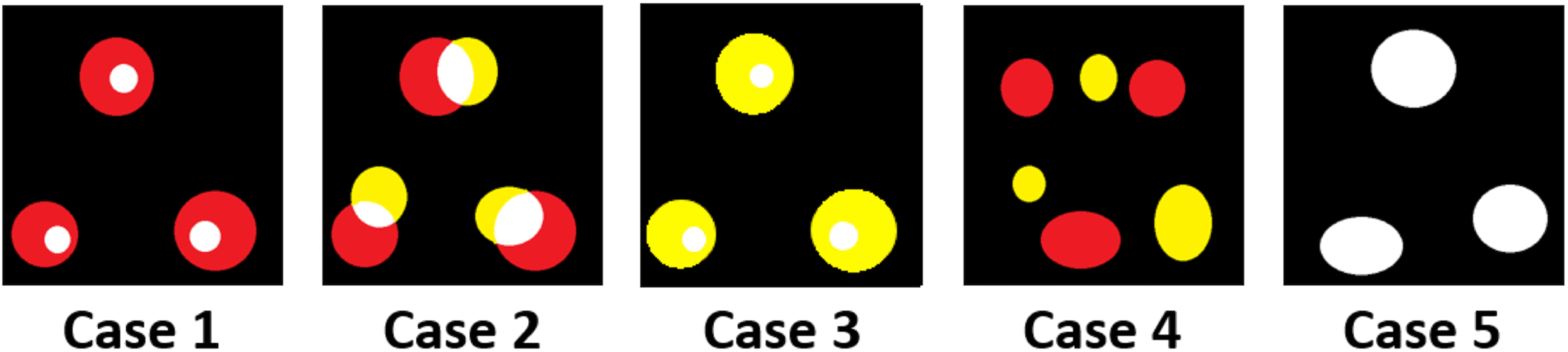
Sample cases for the overall overlap index calculation

**Table 1.**
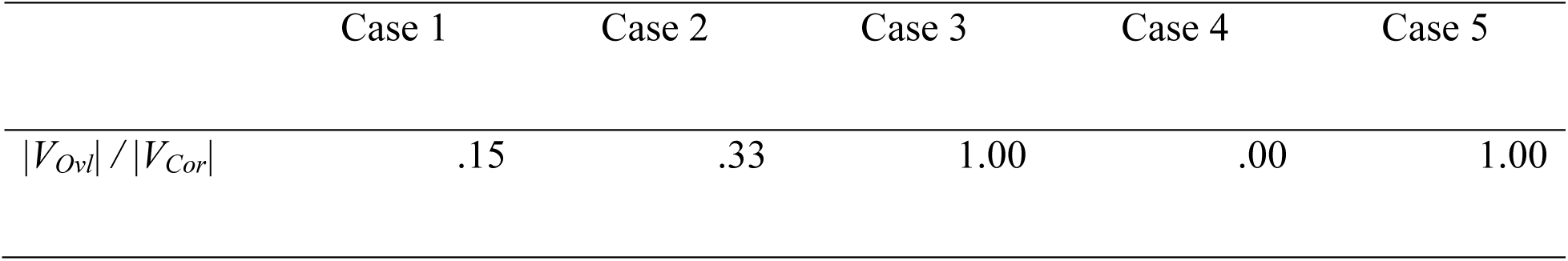
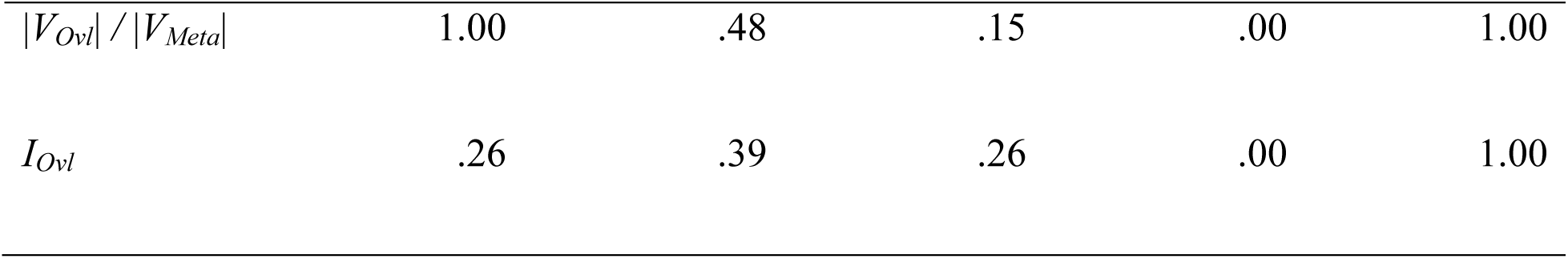
Sole Ratios Compared to Overlap Index for the Cases Depicted in Figure 1

Both case 1 and 3 show why the sole ratio, instead of the overall overlap index, may be misleading when we attempt to evaluate the overlap quantitatively. Although one of two different types of overlap ratios is 1.00, another ratio is significantly smaller and vice versa (.15); in this case, we are not able to decide which one would better represent the overall trend of overlap. Instead, the overall overlap index would be a non-biased solution to address this issue. It can provide us with a unified value for overlap index by taking into account two different types of ratios at the same time. For instance, case 1 and 3 in fact show an identical degree of overlap as visualized in Figure 1 and their overall overlap indices are identical to each other. Furthermore, the overall overlap index value of case 2 is greater than the calculated index values of case 1 and 3; this result is consistent with what we would expect from the apparent ratio of white areas to other areas as presented in the diagrams.

For the evaluation of correction methods in the present study, we calculated the overall overlap index for one contrast, i.e., all moral versus non-moral, because included task conditions in the meta-analysis that intended to be performed are moral cognition and emotion in general, and moral judgment that did not distinguish task conditions according to the nature of moral dilemmas, i.e., moral-personal and moral-impersonal dilemmas. The overlap index was also calculated for case of the overlap between survived voxels after the application of each thresholding method and SnPM. As a result, four overlap indices were calculated (*I*_*Ovl(FDR)*_, *I*_*Ovl(CLU)*_, *I*_*Ovl(RFT)*_, and *I*_*Ovl(Bon)*_) for each type of meta-analysis (either moral cognition and emotion in general, or moral judgment) and SnPM. We examined which correction method produced the highest overlap index value.

## Results

### Reanalysis of fMRI data

We reanalyzed the fMRI to examine which voxels were significantly activated by the different contrasts in the moral judgment task after the application of each correction method. Although we used the spiral in-and-out method that is more robust against the signal dropout in ventral areas in the prefrontal cortex compared to the EPI methods (Glover, 2012; Glover & Law, 2001), some prefrontal regions below z = −12 demonstrated signal loss and were excluded for our reanalysis (see Figure S1). Figure 2 demonstrates survived voxels for each contrast (all moral versus non-moral, moral-personal versus non-moral, moral-impersonal versus non-moral, moral-personal versus moral-impersonal, and moral-impersonal versus moral-personal). Table 2 summarizes the number of survived voxels for each correction method. In addition, Table S2 provides information regarding survived voxels for each contrast. As previous methodological studies have shown, the Bonferroni method-applied FWE voxel-wise thresholding was most conservative, and the FDR-applied thresholding was most lenient (Bennett, Wolford, et al., 2009; Nichols, 2013) among four different thresholding methods in terms of the number of survived voxels (*V*_*FDR*_ ⊃ *V*_*CLU*_ ⊃ *V*_*RFT*_ ⊃ *V*_*Bon*_) for all contrasts, except for the contrast of moral-personal vs. moral-impersonal.

**Figure 2.**
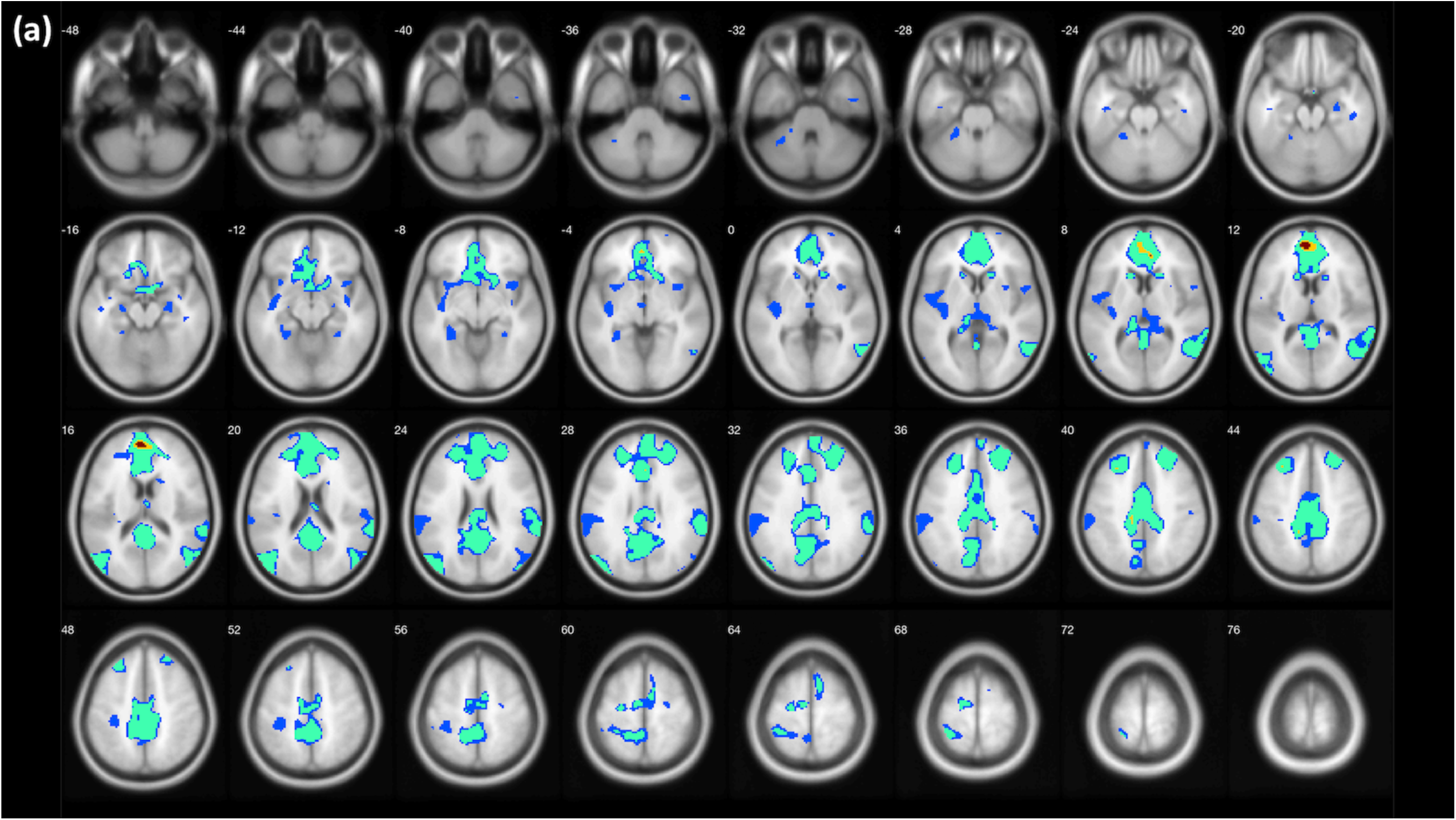
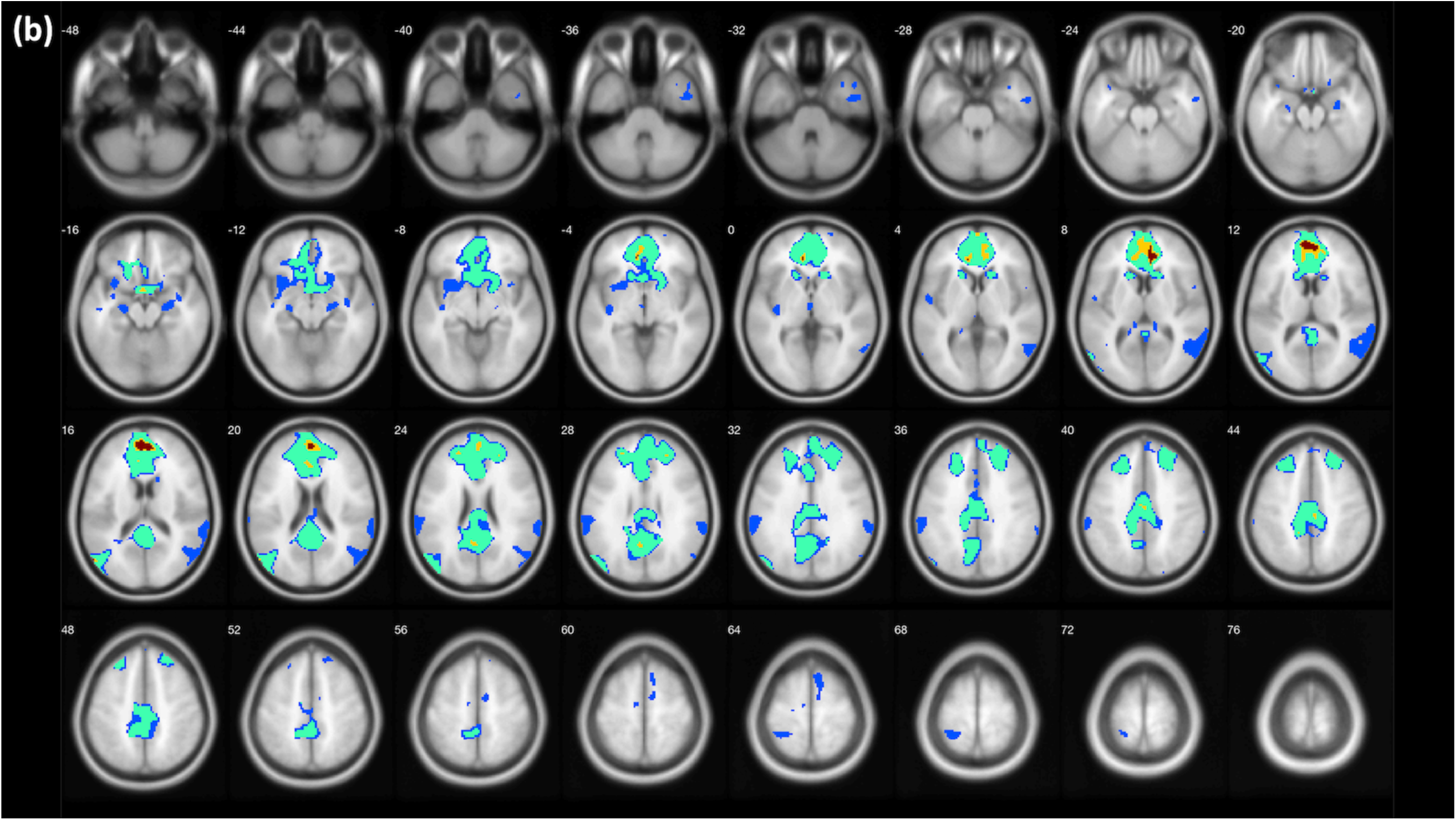
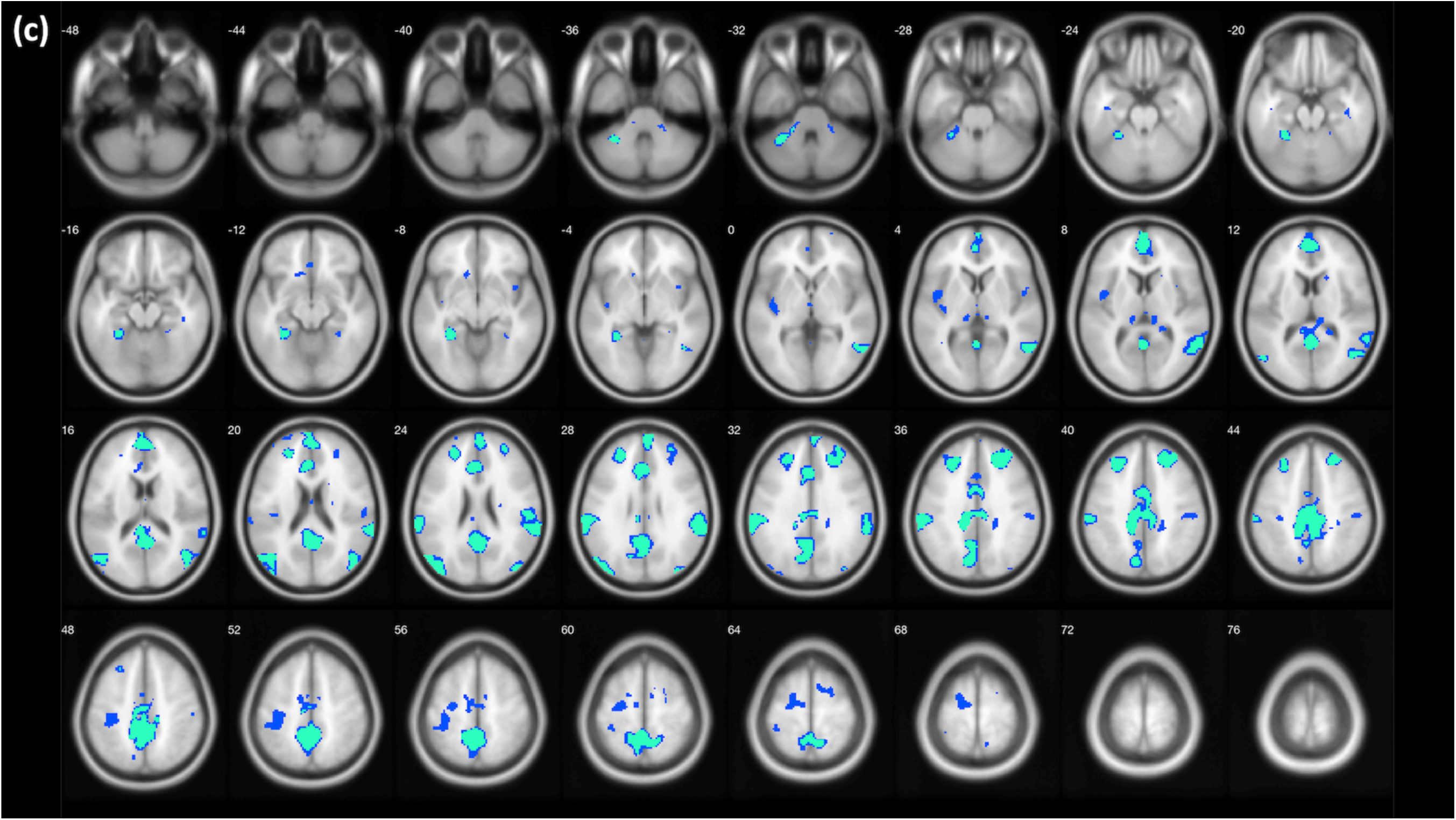
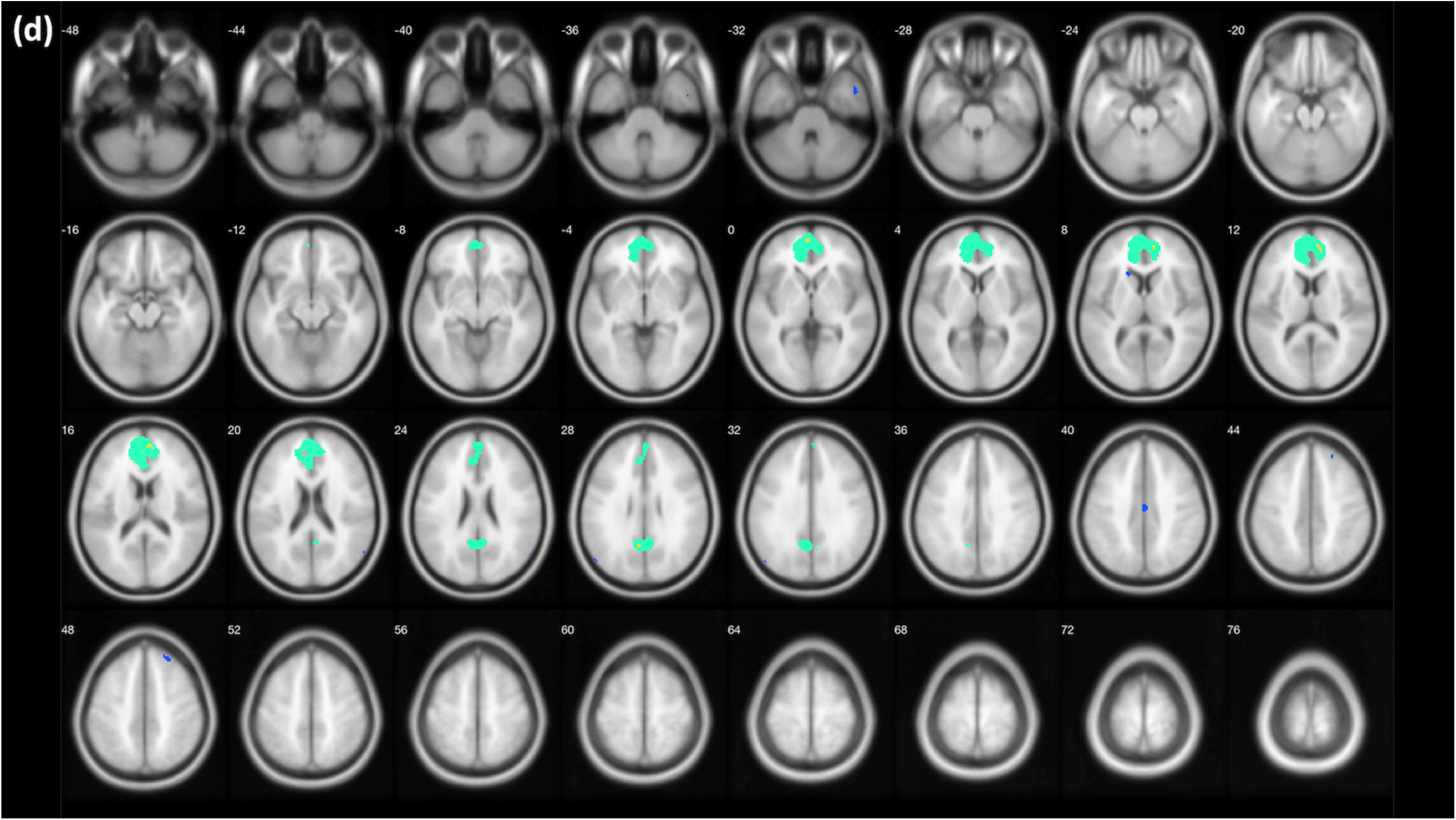
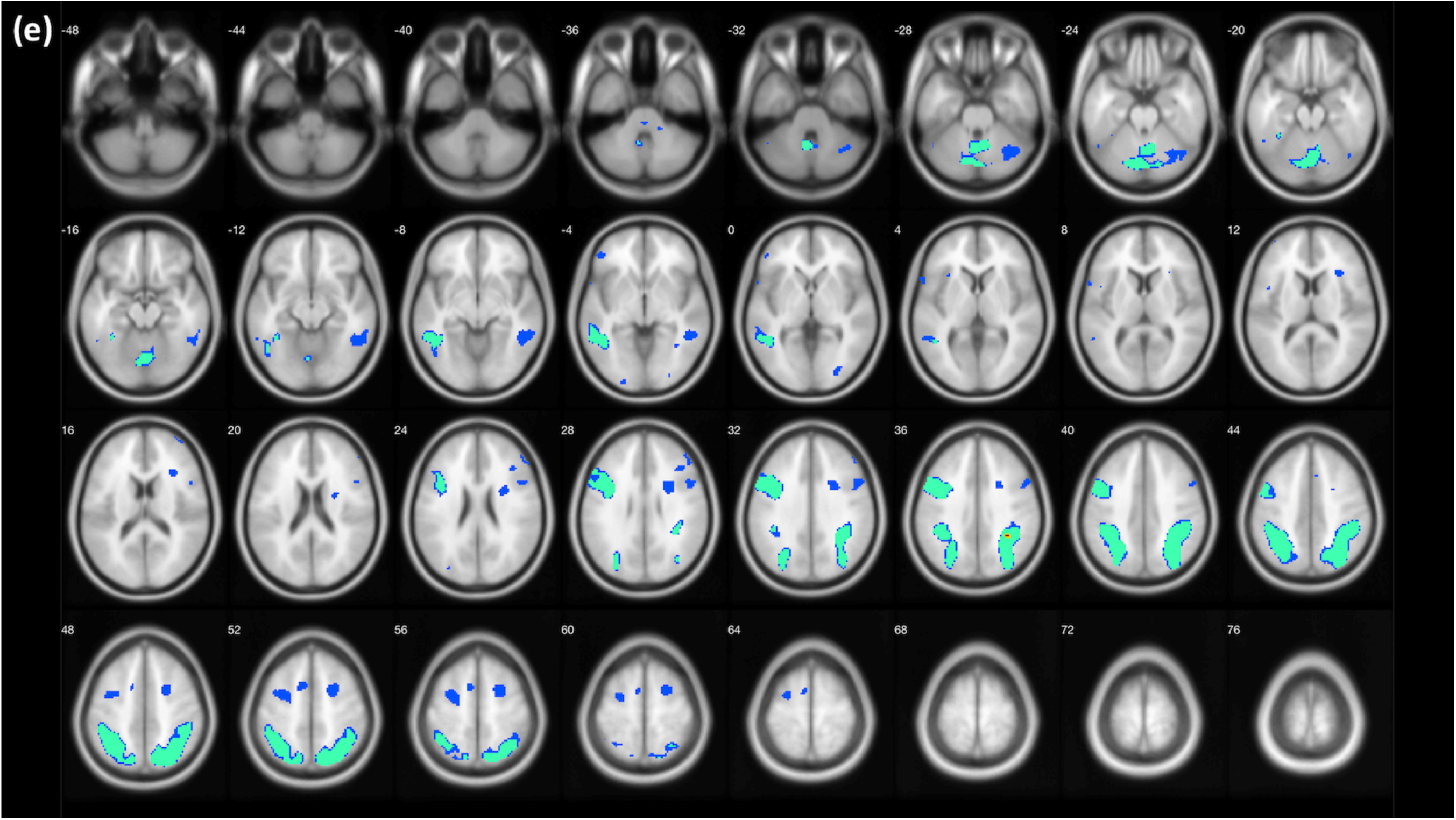
Survived voxels after the application of correction methods. Blue: voxel-wise FDR; green: clusterwise inference; yellow: voxel-wise RFT FWE; red: voxel-wise Bonferroni FWE. (a) All moral (moral-personal + moral-impersonal) vs. non-moral. (b) Moral-personal vs. non-moral. (c) Moral-impersonal vs. non-moral. (d) Moral-personal vs. moral-impersonal. (e) Moral-impersonal vs. moral-personal. Figures created with XjView (Cui, Li, & Song, 2015).

**Table 2.** Number of Voxels Surviving with Four Different Thresholds and SnPM Method as a Reference

| | FDR<br>($\vert V_{FDR}\vert $) | Clusterwise<br>inference<br>($\vert V_{CLU}\vert $) | RFT FWE<br>(voxel-wise)<br>($\vert V_{RFT}\vert $) | Bonferroni's<br>FWE ($\vert V_{Bon}\vert $) | Voxel-wise<br>SnPM<br>($\vert V_{SnPM}\vert $) |
| --- | --- | --- | --- | --- | --- |
| All moral vs. non-moral | 24,524 | 14,424 | 264 | 66 | 416 |
| Moral-personal vs.<br>non-moral | 20,883 | 13,540 | 817 | 227 | 2,084 |
| Moral-impersonal vs.<br>non-moral | 13,787 | 7,458 | 0 | 0 | 20 |
| Moral-personal vs.<br>moral-impersonal | 2,816 | 2,941 | 46 | 0 | 370 |
| Moral-impersonal vs.<br>moral-personal | 11,474 | 6,826 | 22 | 2 | 235 |

### Examination of Overlap with Meta-analysis and SnPM Results

By using the equation for the calculation of the overall overlap index, we examined the quality of each correction method. Survived voxels after the application of each correction method were compared with activation foci identified in the ALE maps created by GingerALE and SnPM. Although some ventral parts of the prefrontal cortex below z = −12 were excluded from our fMRI reanalysis due to signal dropout, no excluded voxels overlapped with any significant voxels in the ALE maps as significant voxels were located above *z* = −8 in the ventromedial prefrontal cortex. Figure 3 demonstrates comparisons with common activation foci of moral cognition and emotion in general, and moral judgment, respectively. Table 3 shows the calculated overall overlap index for each case. As presented, the best overall overlap was achieved when the RFT FWE corrected voxel-wise thresholding was applied. In all cases, the RFT FWE showed the best performance. In the comparisons with the meta-analysis of moral cognition and emotion in general, FDR resulted in 21.4% less overlap with meta-analysis results than RFT FWE, thresholding with clusterwise inference resulted in 2.9% less overlap, and Bonferroni FWE resulted in 66% less overlap. In the comparisons with the meta-analysis of moral judgment, FDR resulted in 72.8% less overlap with meta-analysis results than RFT FWE, thresholding with clusterwise inference resulted in 66.2% less overlap, and Bonferroni FWE resulted in 54.3% less overlap. Finally, when the results were compared with SnPM, FDR resulted in 87.3% less overlap with SnPM results than RFT FWE, thresholding with clusterwise inference resulted in 78.4% less overlap, and Bonferroni FWE resulted in 32.8% less overlap.

**Figure 3.**
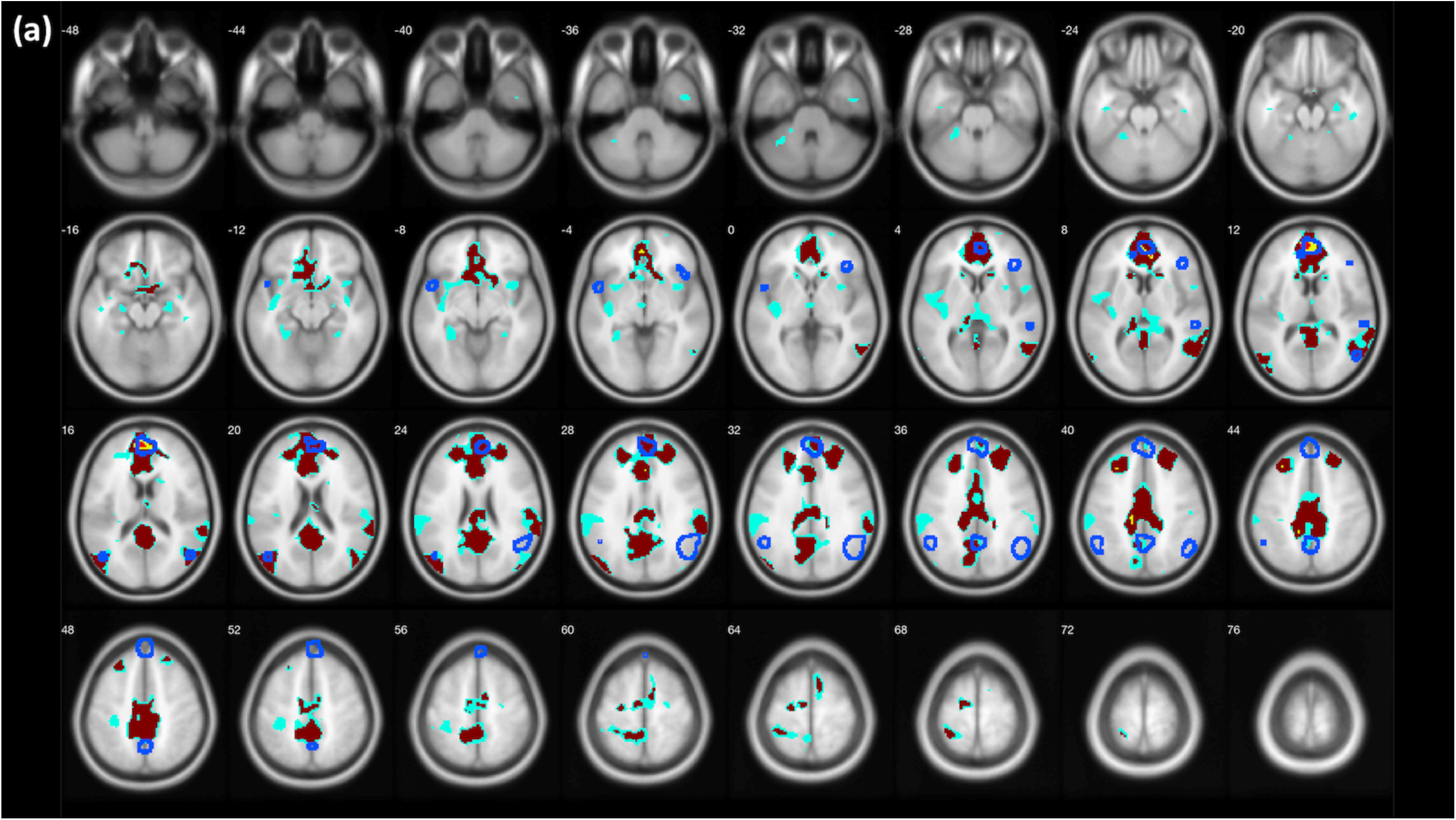
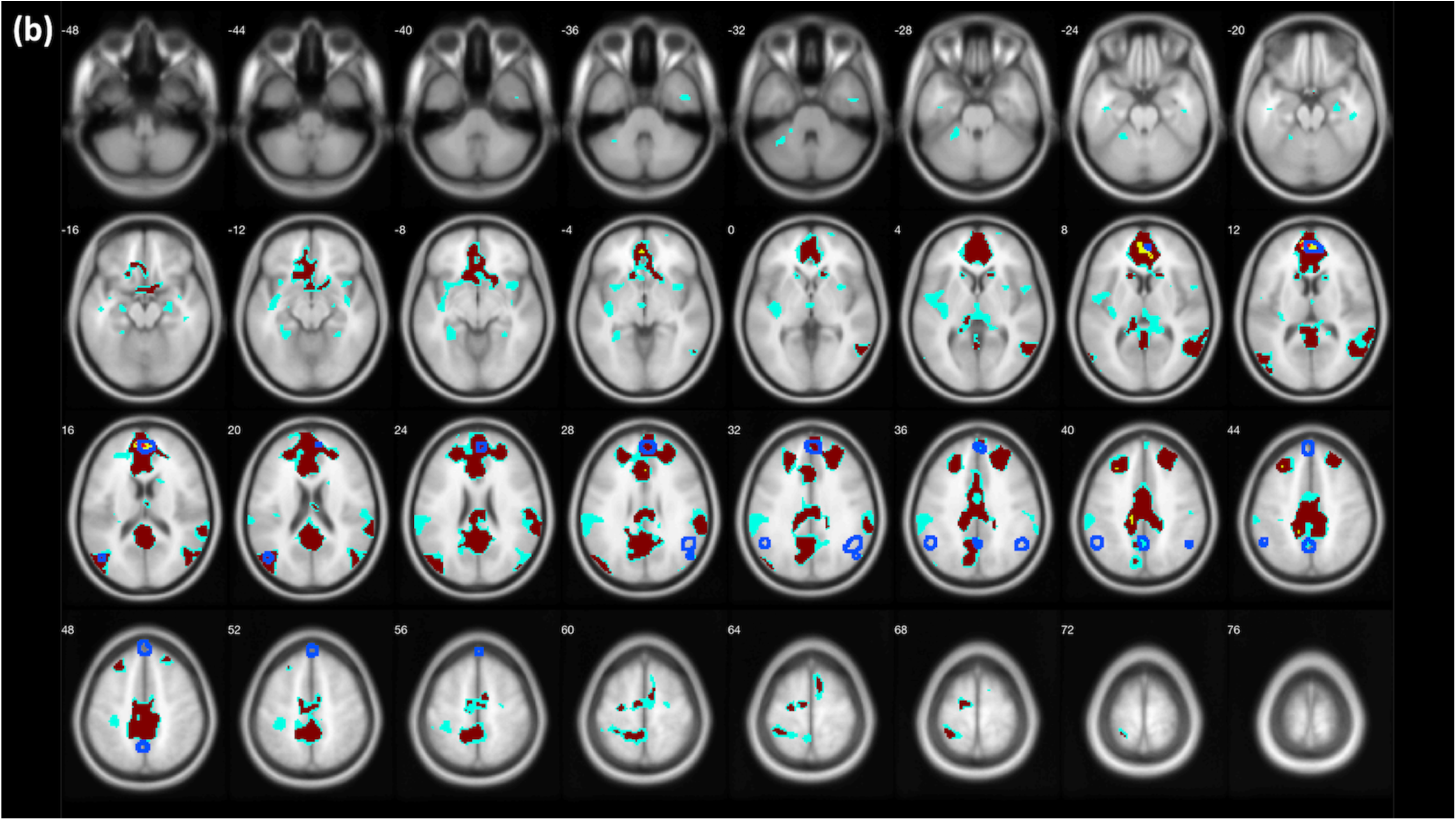
Comparisons between voxels identified using the four correction methods (sky blue: voxel-wise FDR; crimson red: clusterwise inference; yellow: voxel-wise RFT FWE; bright red: voxel-wise Bonferroni FWE) and voxels identified in the meta-analyses and SnPM (areas surrounded by blue lines). (a) Comparisons with meta-analysis of general moral cognition and emotion in general. (b) Comparisons with meta-analysis of moral judgment. (c) Comparisons with voxels identified by SnPM. Figures created with XjView (Cui et al., 2015).

**Table 3.**
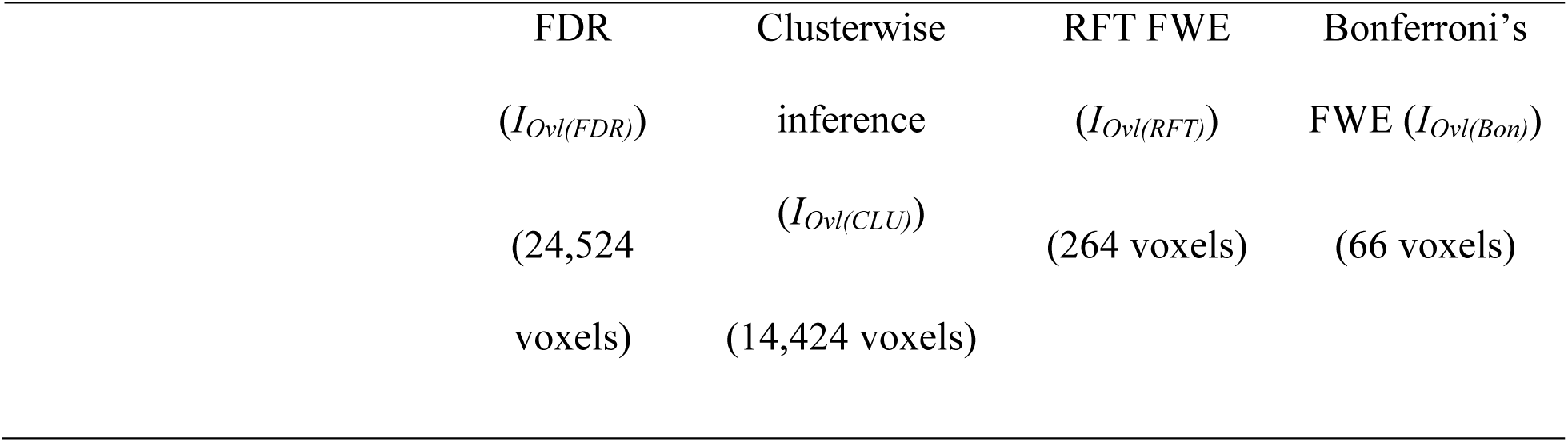
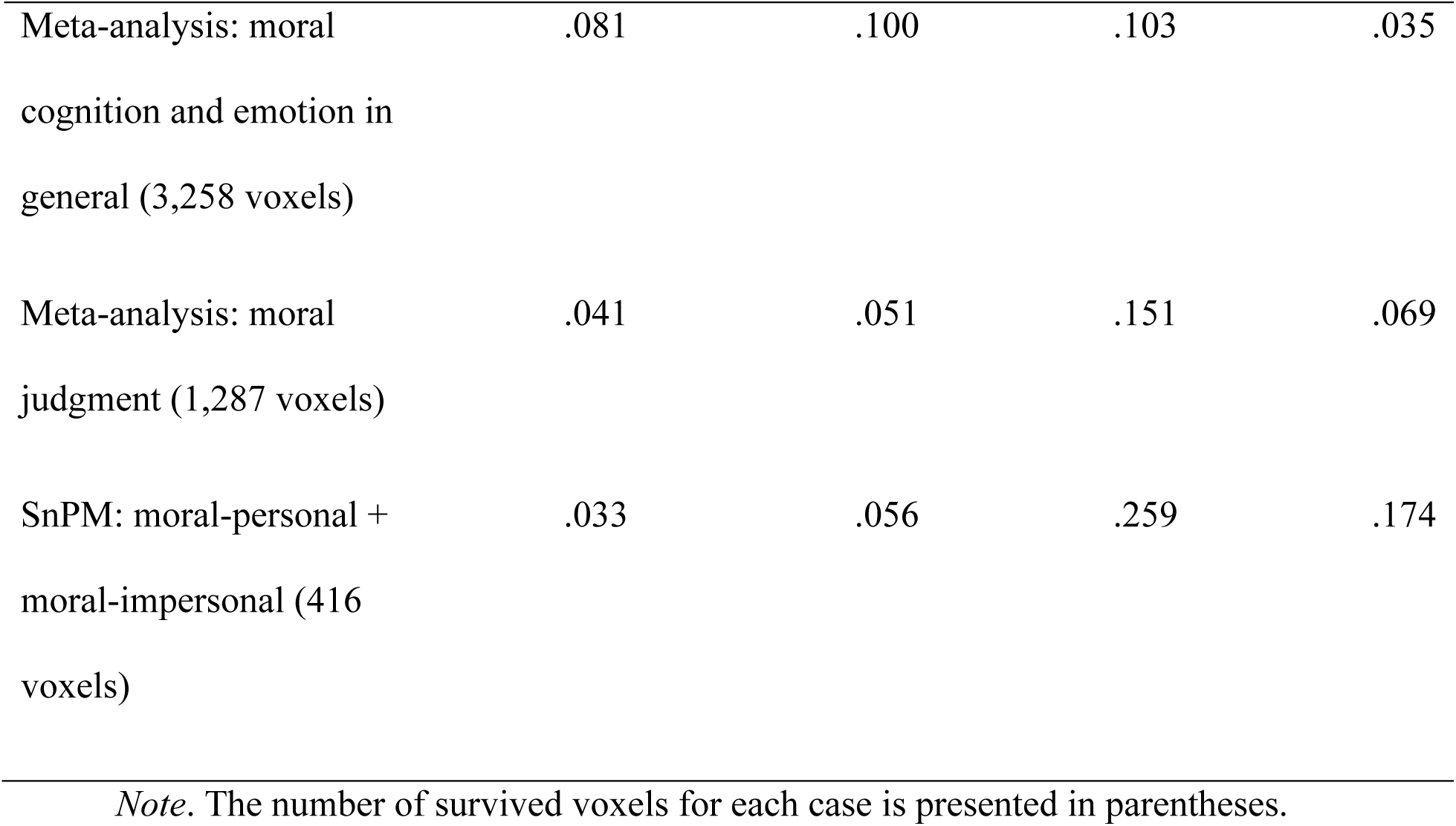
Overlap Index Representing the Degree of Overlap between Activated Voxels Using Each Correction Method and Results from the Meta-Analyses and SnPM

## Discussion

There has been much discussion in the literature about the importance of finding a balance between Type I and Type II errors in fMRI studies. Although some have advocated for less stringent thresholds in order to reduce the risk of missing true effects in social and affective neuroscience studies (Lieberman & Cunningham, 2009), recent studies by Eklund et al. (2016) and Bennett et al. (2009) suggest that some of these methods may be far more lenient than expected or desired. Determining an appropriate method for balancing Type I and Type II error is particularly difficult in social neuroscience studies, which often (though not always) involve complex paradigms that measure less precise and more variable mental processes, resulting in smaller effect sizes and weaker power.

In the present study, we evaluated four different methods for correcting for multiple comparisons within the social neuroscience domain of moral judgment by examining which method produced results that most closely matched the effects identified in a meta-analysis (i.e., an estimate of the “real” effects) and SnPM (i.e., an estimate without any parametric assumptions). As in previous studies, we found that the RFT-applied FWE correction implemented in SPM was similar to or slightly less conservative than the Bonferroni FWE correction method, but more conservative than the FDR correction (Bennett, Wolford, et al., 2009; Eklund et al., 2016; Nichols, 2013; Nichols & Hayasaka, 2003; Nichols & Holmes, 2002) in the case of voxel-wise inference. Also, as recently reported, the clusterwise thresholding provided in SPM12 by default was also more lenient than the voxel-wise FWE-applied thresholding (Eklund et al., 2016). When compared to results from meta-analyses of studies on moral cognition and emotion in general, and moral judgment specifically, as well as those from the application of SnPM, activation maps from the RFT FWE-applied voxel-wise thresholding demonstrated the most overlap, as calculated by the overall overlap index.

These findings suggest that the RFT FWE-applied voxel-wise thresholding may be an acceptable correction method for studies of moral psychology despite the limitations that have been discussed in the field of social neuroscience (e.g., conservativeness and a lack of statistical power). The RFT FWE-applied voxel-wise thresholding may provide a more appropriate balance between false positives and false negatives than other correction methods. Although this method implemented in SPM is susceptible to reduced sensitivity, in the context of some areas of social neuroscience the RFT FWE-applied voxel-wise thresholding may still be considered a viable correction method given the calculated overall overlap indices despite the statistical power issue. However, researchers in social neuroscience should seriously consider cross-checking the thresholding method with meta-analyses and/or SnPM in order to evaluate whether it is appropriate to apply the method in the context of their study.

In the present study, we compared different correction methods in the context of a moral judgment task. However, this unique approach of comparing the results from different correction methods to those of a meta-analysis could be applied to many different domains of social and affective neuroscience. Future studies using this approach can provide more information about whether voxel-wise RFT FWE results in the most overlap with meta-analysis results in other domains within social and affective neuroscience.

The primary limitation of the present study is the imperfect nature of the meta-analysis that was used to examine the quality of the multiple correction methods. The meta-analysis was based on studies that implement a variety of tasks related to moral cognition and emotion, and that use a variety of correction methods and thresholds, thus introducing the possibility of biased results. However, by aggregating data from many studies, the hope is that the meta-analysis technique can provide a relatively unbiased indicator of the areas that appear to be commonly activated across many studies – particularly in social neuroscience studies that involve rather complex processes. In addition, as Eklund et al. (2016) warned that the employment of meta-analysis does not necessarily mitigate the need for the application of valid inferential methodologies for individual studies.

Another limitation is that, as is common in meta-analyses, our meta-analysis was not limited to previous studies that utilized the same paradigm (Greene et al.’s (2001, 2004) experimental design). Thus, the activation foci identified in the meta-analysis might include brain voxels not directly associated with moral judgment, the process of interest, and might be inappropriate to be used for a comparison and evaluation. Of course, the best way to address this issue is to only meta-analyze published studies that used Greene et al.’s (2001, 2004) dilemmas; however, due to the limited number of studies, this was not feasible. To address this issue, we tried to apply a more stringent inclusion rule (i.e., meta-analyzing studies related to moral judgment instead of moral cognition and emotion in general) in order to address this issue; the results demonstrated that there were greater overlaps between activation foci found by the aforementioned fMRI experiment and activation foci found by a meta-analysis of studies related moral judgment compared to those found by a meta-analysis of studies related moral cognition and emotion in general.

Given these limitations associated with meta-analysis, researchers may have to employ additional cross-checking methods, such as SnPM, which is free from any error originating from parametric assumptions, to provide further evaluation of findings (Nichols & Hayasaka, 2003). Researchers may also consider utilizing alternative correction methods not assessed here, such as the threshold-free cluster enhancement (TFCE), which is considered to have greater sensitivity than the traditional methods and allows for the false positive rate to be set at a predetermined level by the permutation test (Smith & Nichols, 2009). Although, this function has not been implemented in SPM, which was examined in the present study, it is available in FSL as an option in the randomise permutation-based inference tool.

## Conclusion

The goal of correcting for multiple comparisons in fMRI studies is to generate clusters of activity that reflect true effects, and thus would be expected to replicate in future studies. Here we show that using the RFT FWE-applied voxel-wise thresholding method in a study of moral judgment produced the most overlap with results from a meta-analysis on moral cognition and moral emotion and the most overlap with SnPM analyses, suggesting that this method may be the best for achieving the goal of identifying true effects. Although this method suffers from potentially insufficient statistical power, which has been a significant issue in social neuroscience, it may be an acceptable option in the context of experiments focusing on morality and possibly other domains of social neuroscience, as long as its application is cross-checked with other methods.

## Supplementary Materials

**Table S1.**
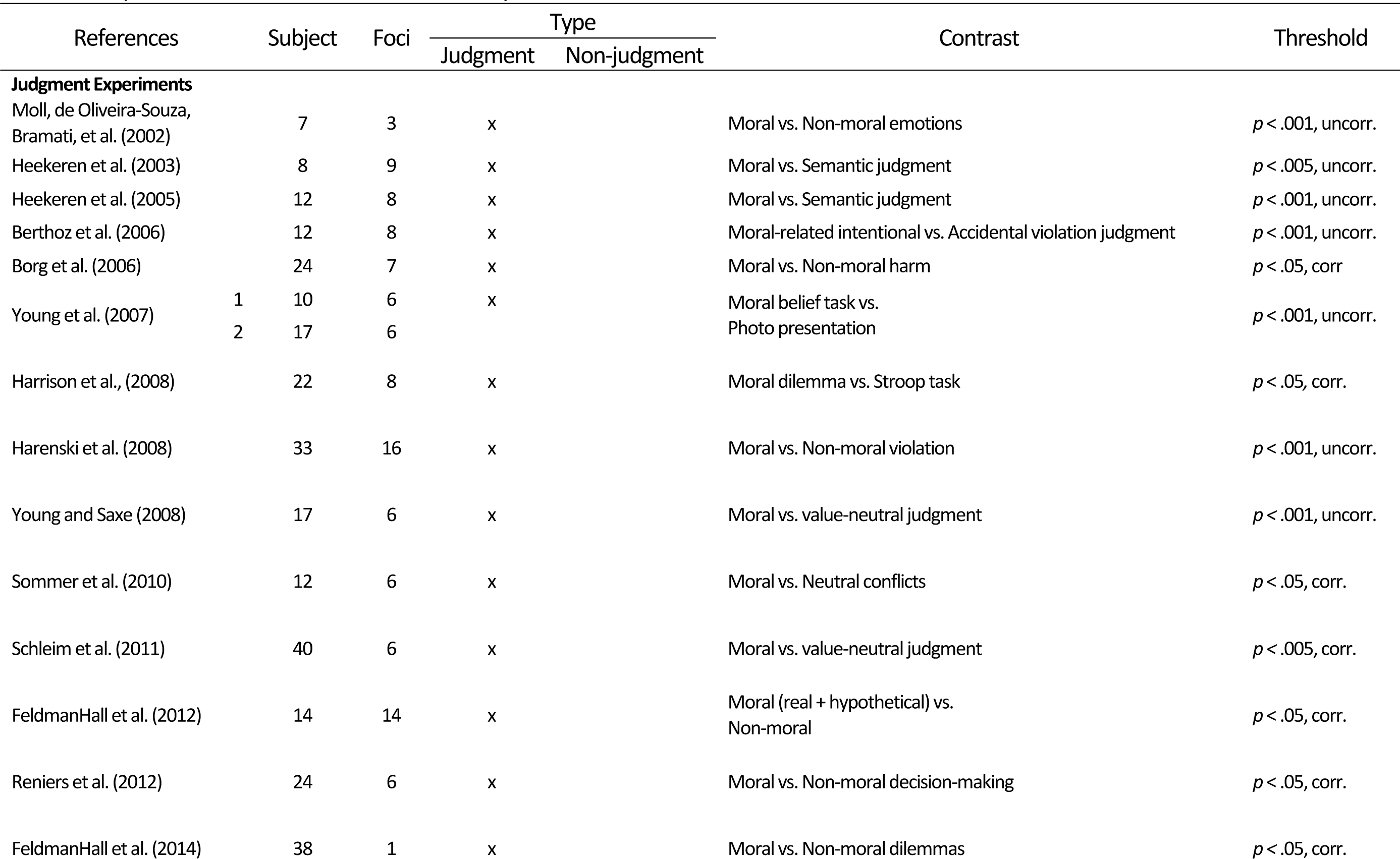
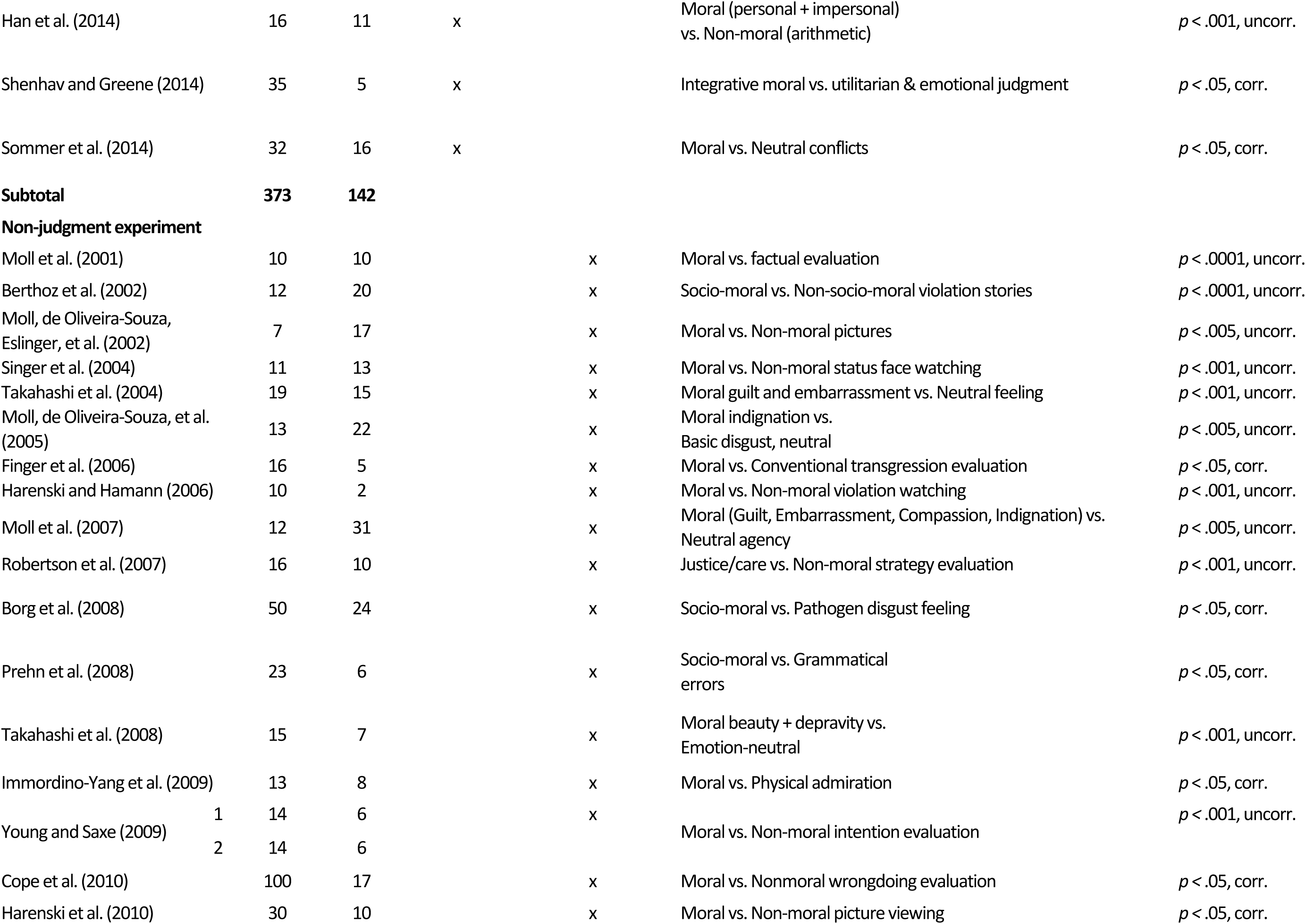
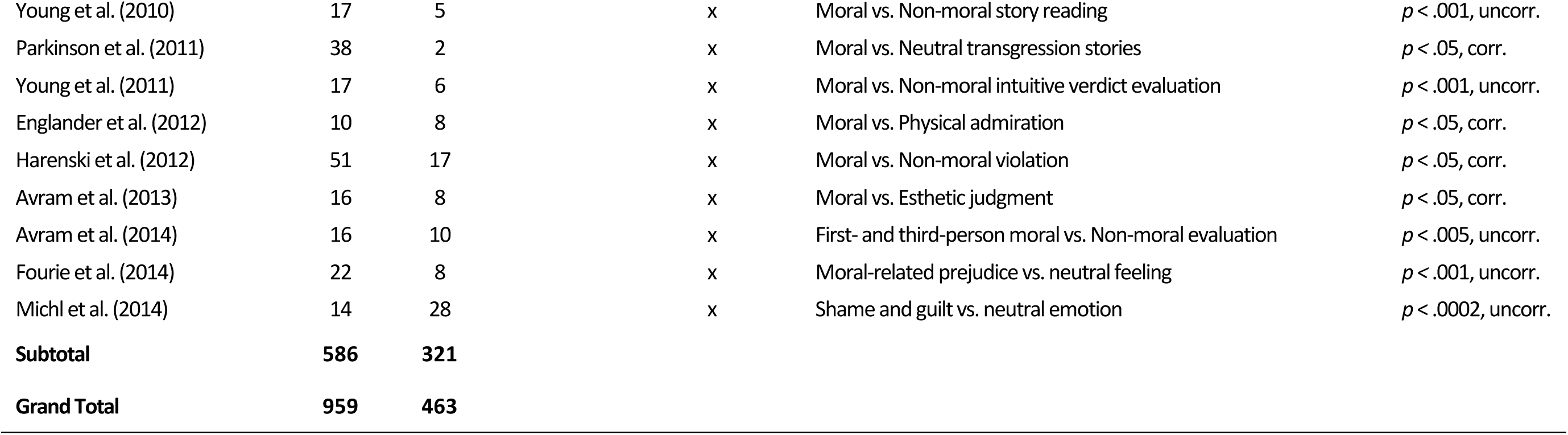
Experiments Included in the Meta-analysis

---

**Table S2.**
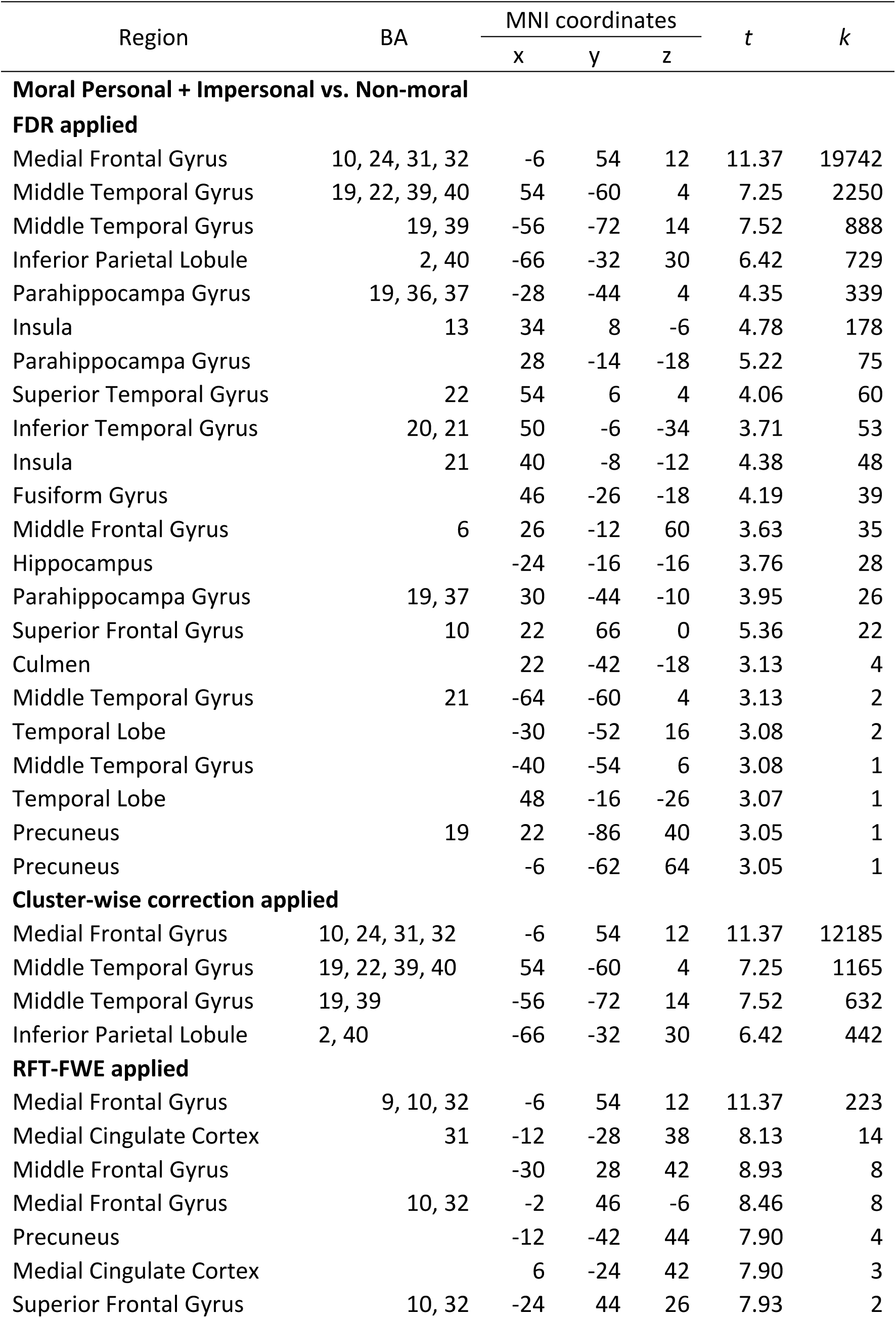
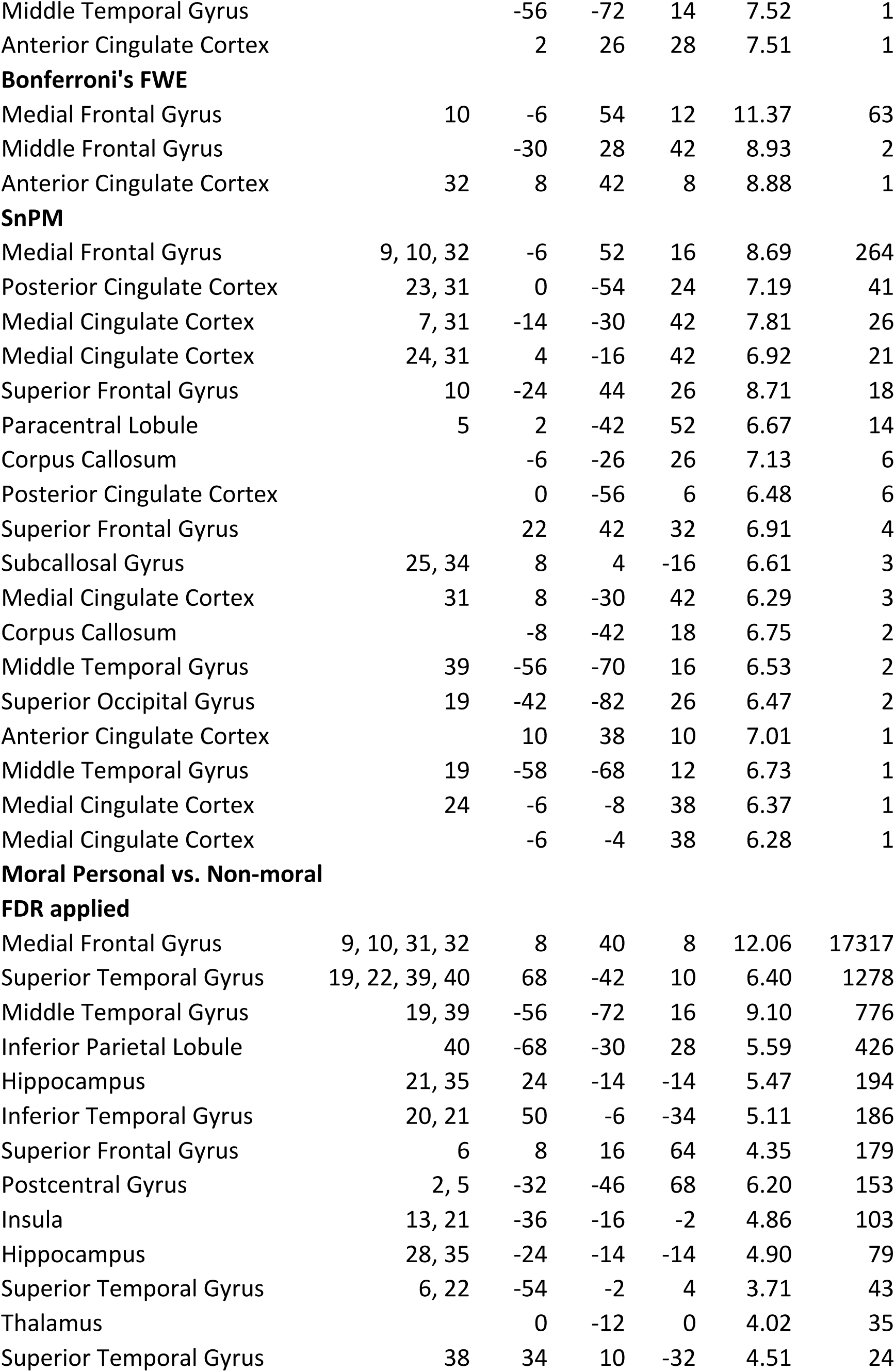
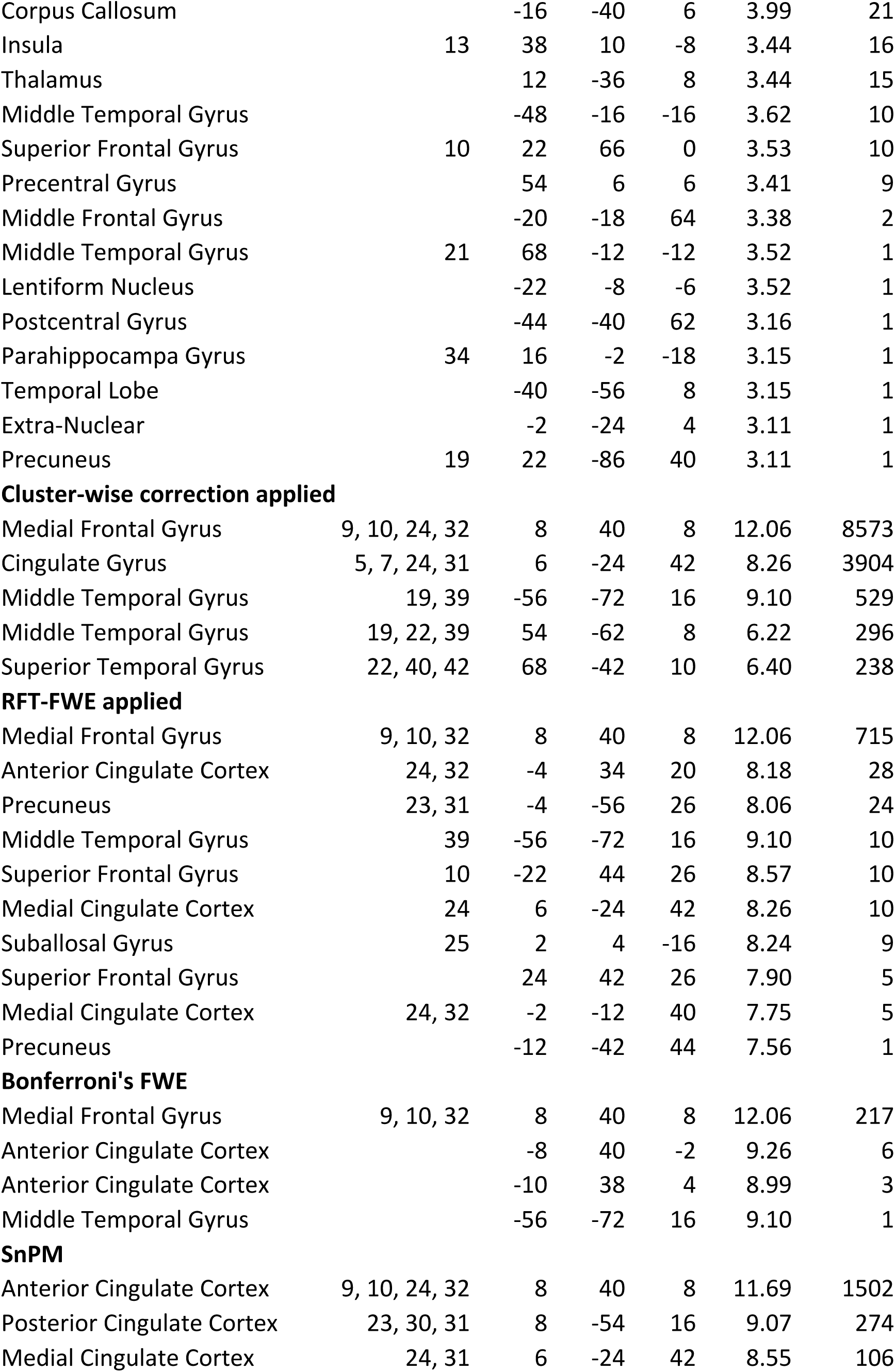
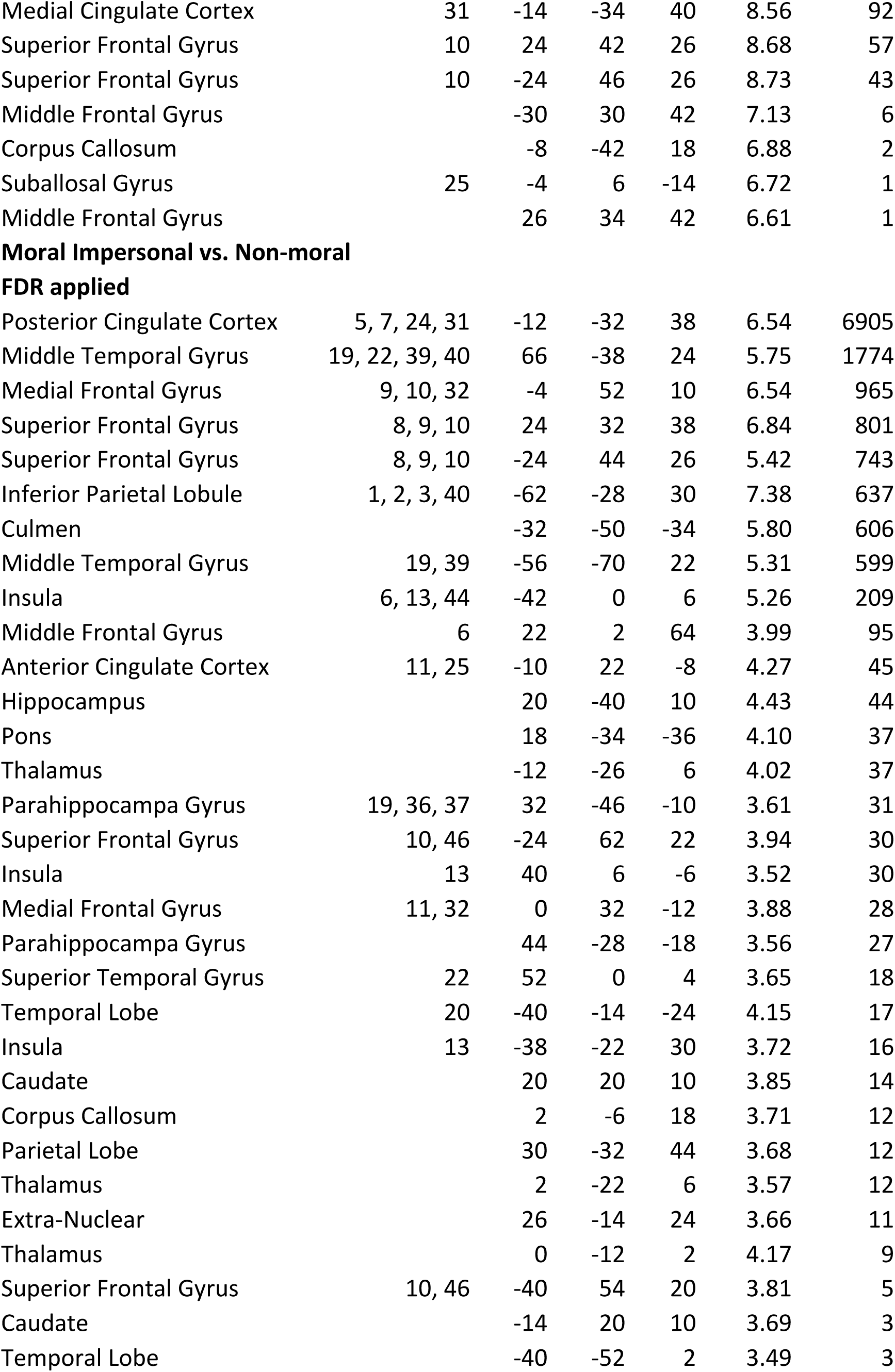
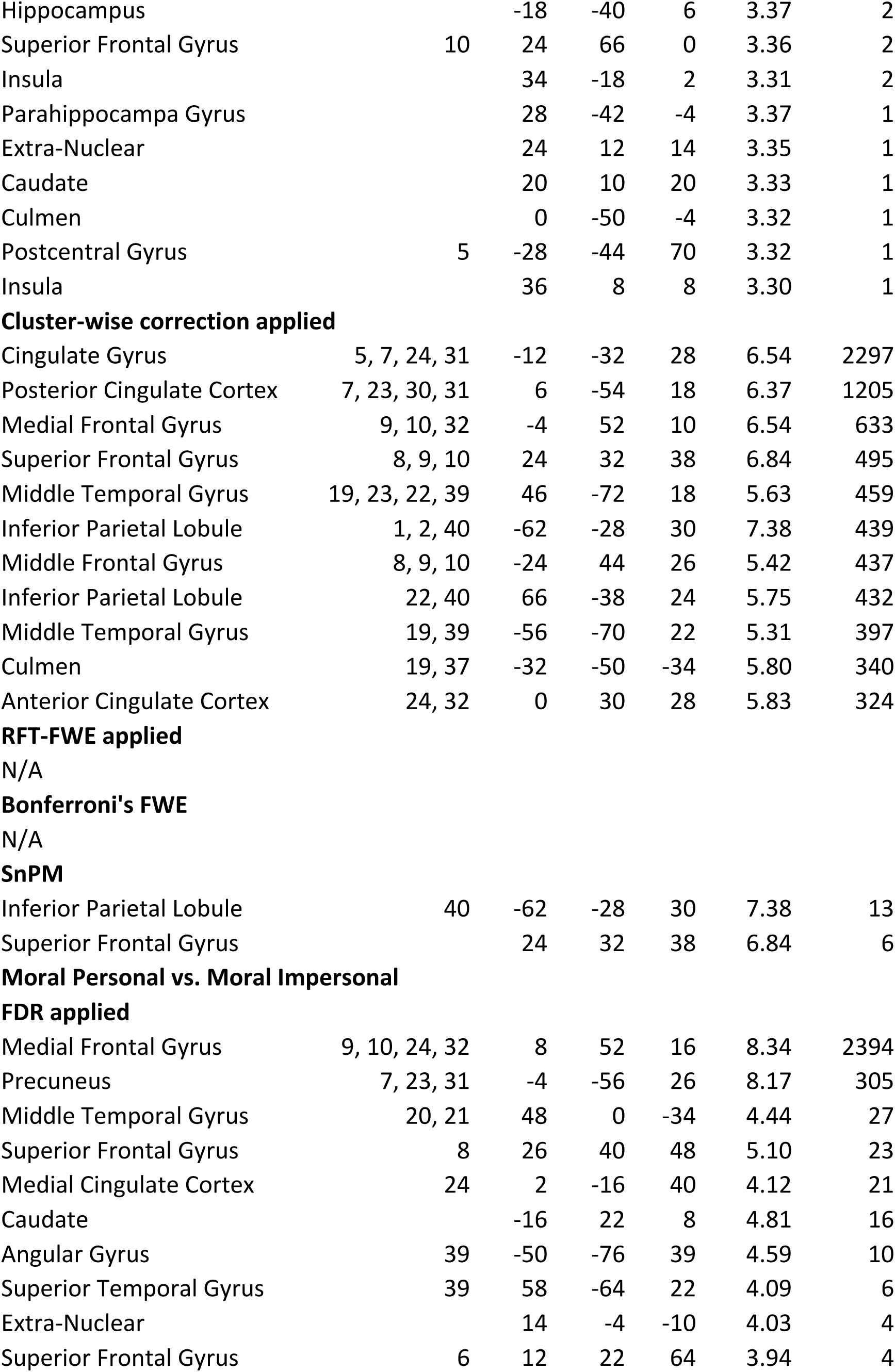
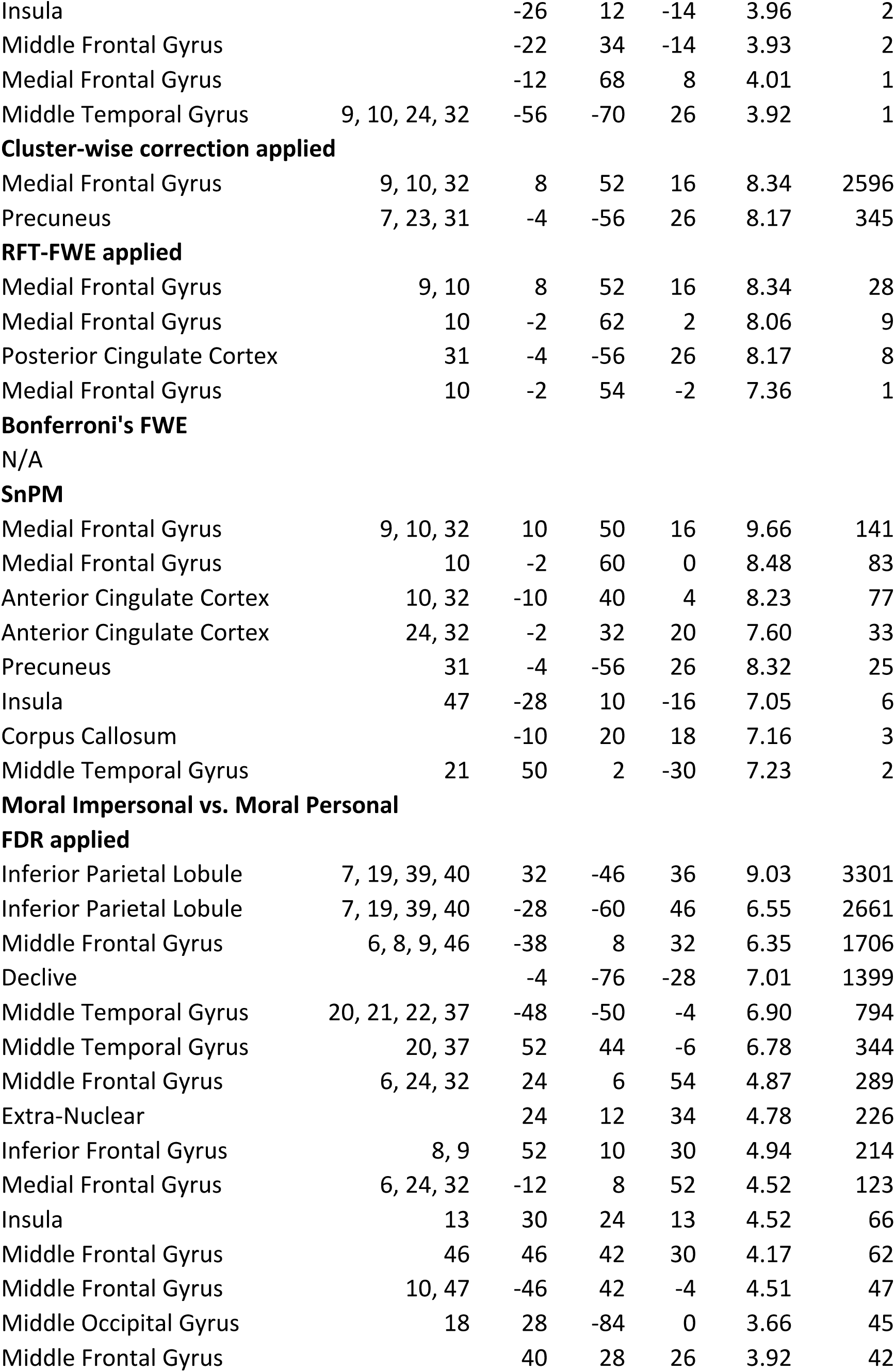
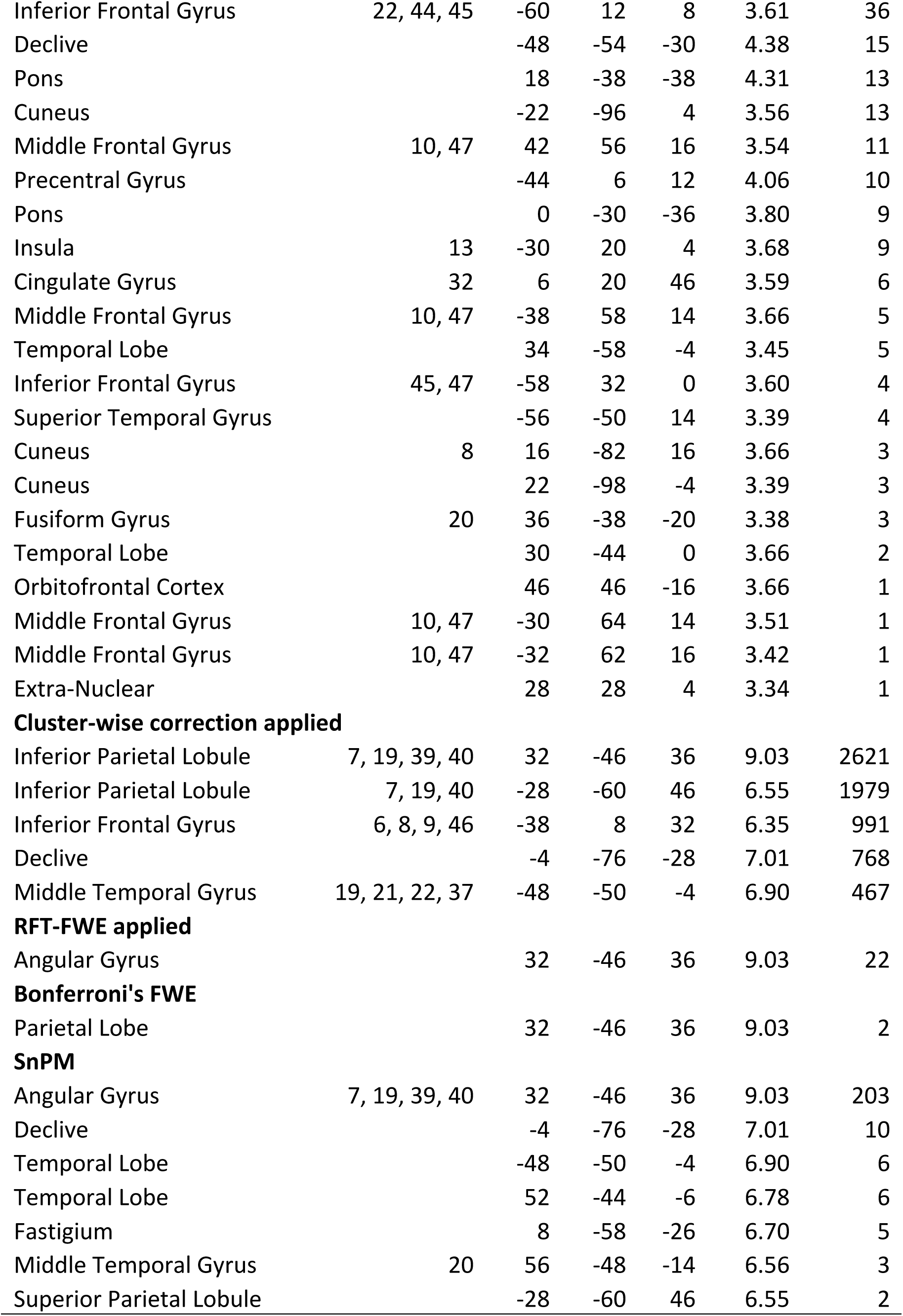
Activation foci for each contrast

| Region | BA | MNI coordinates |  |  | t | k |
| --- | --- | --- | --- | --- | --- | --- |
|  |  | x | y | z |  |  |
| Moral Personal + Impersonal vs. Non-moral |  |  |  |  |  |  |
| FDR applied |  |  |  |  |  |  |
| Medial Frontal Gyrus | 10, 24, 31, 32 | -6 | 54 | 12 | 11.37 | 19742 |
| Middle Temporal Gyrus | 19, 22, 39, 40 | 54 | -60 | 4 | 7.25 | 2250 |
| Middle Temporal Gyrus | 19, 39 | -56 | -72 | 14 | 7.52 | 888 |
| Inferior Parietal Lobule | 2, 40 | -66 | -32 | 30 | 6.42 | 729 |
| Parahippocampa Gyrus | 19, 36, 37 | -28 | -44 | 4 | 4.35 | 339 |
| Insula | 13 | 34 | 8 | -6 | 4.78 | 178 |
| Parahippocampa Gyrus |  | 28 | -14 | -18 | 5.22 | 75 |
| Superior Temporal Gyrus | 22 | 54 | 6 | 4 | 4.06 | 60 |
| Inferior Temporal Gyrus | 20, 21 | 50 | -6 | -34 | 3.71 | 53 |
| Insula | 21 | 40 | -8 | -12 | 4.38 | 48 |
| Fusiform Gyrus |  | 46 | -26 | -18 | 4.19 | 39 |
| Middle Frontal Gyrus | 6 | 26 | -12 | 60 | 3.63 | 35 |
| Hippocampus |  | -24 | -16 | -16 | 3.76 | 28 |
| Parahippocampa Gyrus | 19, 37 | 30 | -44 | -10 | 3.95 | 26 |
| Superior Frontal Gyrus | 10 | 22 | 66 | 0 | 5.36 | 22 |
| Culmen |  | 22 | -42 | -18 | 3.13 | 4 |
| Middle Temporal Gyrus | 21 | -64 | -60 | 4 | 3.13 | 2 |
| Temporal Lobe |  | -30 | -52 | 16 | 3.08 | 2 |
| Middle Temporal Gyrus |  | -40 | -54 | 6 | 3.08 | 1 |
| Temporal Lobe |  | 48 | -16 | -26 | 3.07 | 1 |
| Precuneus | 19 | 22 | -86 | 40 | 3.05 | 1 |
| Precuneus |  | -6 | -62 | 64 | 3.05 | 1 |
| Cluster-wise correction applied |  |  |  |  |  |  |
| Medial Frontal Gyrus | 10, 24, 31, 32 | -6 | 54 | 12 | 11.37 | 12185 |
| Middle Temporal Gyrus | 19, 22, 39, 40 | 54 | -60 | 4 | 7.25 | 1165 |
| Middle Temporal Gyrus | 19, 39 | -56 | -72 | 14 | 7.52 | 632 |
| Inferior Parietal Lobule | 2, 40 | -66 | -32 | 30 | 6.42 | 442 |
| RFT-FWE applied |  |  |  |  |  |  |
| Medial Frontal Gyrus | 9, 10, 32 | -6 | 54 | 12 | 11.37 | 223 |
| Medial Cingulate Cortex | 31 | -12 | -28 | 38 | 8.13 | 14 |
| Middle Frontal Gyrus |  | -30 | 28 | 42 | 8.93 | 8 |
| Medial Frontal Gyrus | 10, 32 | -2 | 46 | -6 | 8.46 | 8 |
| Precuneus |  | -12 | -42 | 44 | 7.90 | 4 |
| Medial Cingulate Cortex |  | 6 | -24 | 42 | 7.90 | 3 |
| Superior Frontal Gyrus | 10, 32 | -24 | 44 | 26 | 7.93 | 2 |
| Middle Temporal Gyrus |  | -56 | -72 | 14 | 7.52 | 1 |
| Anterior Cingulate Cortex |  | 2 | 26 | 28 | 7.51 | 1 |
| **Bonferroni's FWE** |  |  |  |  |  |  |
| Medial Frontal Gyrus | 10 | -6 | 54 | 12 | 11.37 | 63 |
| Middle Frontal Gyrus |  | -30 | 28 | 42 | 8.93 | 2 |
| Anterior Cingulate Cortex | 32 | 8 | 42 | 8 | 8.88 | 1 |
| **SnPM** |  |  |  |  |  |  |
| Medial Frontal Gyrus | 9, 10, 32 | -6 | 52 | 16 | 8.69 | 264 |
| Posterior Cingulate Cortex | 23, 31 | 0 | -54 | 24 | 7.19 | 41 |
| Medial Cingulate Cortex | 7, 31 | -14 | -30 | 42 | 7.81 | 26 |
| Medial Cingulate Cortex | 24, 31 | 4 | -16 | 42 | 6.92 | 21 |
| Superior Frontal Gyrus | 10 | -24 | 44 | 26 | 8.71 | 18 |
| Paracentral Lobule | 5 | 2 | -42 | 52 | 6.67 | 14 |
| Corpus Callosum |  | -6 | -26 | 26 | 7.13 | 6 |
| Posterior Cingulate Cortex |  | 0 | -56 | 6 | 6.48 | 6 |
| Superior Frontal Gyrus |  | 22 | 42 | 32 | 6.91 | 4 |
| Subcallosal Gyrus | 25, 34 | 8 | 4 | -16 | 6.61 | 3 |
| Medial Cingulate Cortex | 31 | 8 | -30 | 42 | 6.29 | 3 |
| Corpus Callosum |  | -8 | -42 | 18 | 6.75 | 2 |
| Middle Temporal Gyrus | 39 | -56 | -70 | 16 | 6.53 | 2 |
| Superior Occipital Gyrus | 19 | -42 | -82 | 26 | 6.47 | 2 |
| Anterior Cingulate Cortex |  | 10 | 38 | 10 | 7.01 | 1 |
| Middle Temporal Gyrus | 19 | -58 | -68 | 12 | 6.73 | 1 |
| Medial Cingulate Cortex | 24 | -6 | -8 | 38 | 6.37 | 1 |
| Medial Cingulate Cortex |  | -6 | -4 | 38 | 6.28 | 1 |
| **FDR applied** |  |  |  |  |  |  |
| Medial Frontal Gyrus | 9, 10, 31, 32 | 8 | 40 | 8 | 12.06 | 17317 |
| Superior Temporal Gyrus | 19, 22, 39, 40 | 68 | -42 | 10 | 6.40 | 1278 |
| Middle Temporal Gyrus | 19, 39 | -56 | -72 | 16 | 9.10 | 776 |
| Inferior Parietal Lobule | 40 | -68 | -30 | 28 | 5.59 | 426 |
| Hippocampus | 21, 35 | 24 | -14 | -14 | 5.47 | 194 |
| Inferior Temporal Gyrus | 20, 21 | 50 | -6 | -34 | 5.11 | 186 |
| Superior Frontal Gyrus | 6 | 8 | 16 | 64 | 4.35 | 179 |
| Postcentral Gyrus | 2, 5 | -32 | -46 | 68 | 6.20 | 153 |
| Insula | 13, 21 | -36 | -16 | -2 | 4.86 | 103 |
| Hippocampus | 28, 35 | -24 | -14 | -14 | 4.90 | 79 |
| Superior Temporal Gyrus | 6, 22 | -54 | -2 | 4 | 3.71 | 43 |
| Thalamus |  | 0 | -12 | 0 | 4.02 | 35 |
| Superior Temporal Gyrus | 38 | 34 | 10 | -32 | 4.51 | 24 |
| Corpus Callosum |  | -16 | -40 | 6 | 3.99 | 21 |
| Insula | 13 | 38 | 10 | -8 | 3.44 | 16 |
| Thalamus |  | 12 | -36 | 8 | 3.44 | 15 |
| Middle Temporal Gyrus |  | -48 | -16 | -16 | 3.62 | 10 |
| Superior Frontal Gyrus | 10 | 22 | 66 | 0 | 3.53 | 10 |
| Precentral Gyrus |  | 54 | 6 | 6 | 3.41 | 9 |
| Middle Frontal Gyrus |  | -20 | -18 | 64 | 3.38 | 2 |
| Middle Temporal Gyrus | 21 | 68 | -12 | -12 | 3.52 | 1 |
| Lentiform Nucleus |  | -22 | -8 | -6 | 3.52 | 1 |
| Postcentral Gyrus |  | -44 | -40 | 62 | 3.16 | 1 |
| Parahippocampa Gyrus | 34 | 16 | -2 | -18 | 3.15 | 1 |
| Temporal Lobe |  | -40 | -56 | 8 | 3.15 | 1 |
| Extra-Nuclear |  | -2 | -24 | 4 | 3.11 | 1 |
| Precuneus | 19 | 22 | -86 | 40 | 3.11 | 1 |
| **Cluster-wise correction applied** |  |  |  |  |  |  |
| Medial Frontal Gyrus | 9, 10, 24, 32 | 8 | 40 | 8 | 12.06 | 8573 |
| Cingulate Gyrus | 5, 7, 24, 31 | 6 | -24 | 42 | 8.26 | 3904 |
| Middle Temporal Gyrus | 19, 39 | -56 | -72 | 16 | 9.10 | 529 |
| Middle Temporal Gyrus | 19, 22, 39 | 54 | -62 | 8 | 6.22 | 296 |
| Superior Temporal Gyrus | 22, 40, 42 | 68 | -42 | 10 | 6.40 | 238 |
| **RFT-FWE applied** |  |  |  |  |  |  |
| Medial Frontal Gyrus | 9, 10, 32 | 8 | 40 | 8 | 12.06 | 715 |
| Anterior Cingulate Cortex | 24, 32 | -4 | 34 | 20 | 8.18 | 28 |
| Precuneus | 23, 31 | -4 | -56 | 26 | 8.06 | 24 |
| Middle Temporal Gyrus | 39 | -56 | -72 | 16 | 9.10 | 10 |
| Superior Frontal Gyrus | 10 | -22 | 44 | 26 | 8.57 | 10 |
| Medial Cingulate Cortex | 24 | 6 | -24 | 42 | 8.26 | 10 |
| Suballosal Gyrus | 25 | 2 | 4 | -16 | 8.24 | 9 |
| Superior Frontal Gyrus |  | 24 | 42 | 26 | 7.90 | 5 |
| Medial Cingulate Cortex | 24, 32 | -2 | -12 | 40 | 7.75 | 5 |
| Precuneus |  | -12 | -42 | 44 | 7.56 | 1 |
| **Bonferroni's FWE** |  |  |  |  |  |  |
| Medial Frontal Gyrus | 9, 10, 32 | 8 | 40 | 8 | 12.06 | 217 |
| Anterior Cingulate Cortex |  | -8 | 40 | -2 | 9.26 | 6 |
| Anterior Cingulate Cortex |  | -10 | 38 | 4 | 8.99 | 3 |
| Middle Temporal Gyrus |  | -56 | -72 | 16 | 9.10 | 1 |
| **SnPM** |  |  |  |  |  |  |
| Anterior Cingulate Cortex | 9, 10, 24, 32 | 8 | 40 | 8 | 11.69 | 1502 |
| Posterior Cingulate Cortex | 23, 30, 31 | 8 | -54 | 16 | 9.07 | 274 |
| Medial Cingulate Cortex | 24, 31 | 6 | -24 | 42 | 8.55 | 106 |
| Medial Cingulate Cortex | 31 | -14 | -34 | 40 | 8.56 | 92 |
| Superior Frontal Gyrus | 10 | 24 | 42 | 26 | 8.68 | 57 |
| Superior Frontal Gyrus | 10 | -24 | 46 | 26 | 8.73 | 43 |
| Middle Frontal Gyrus |  | -30 | 30 | 42 | 7.13 | 6 |
| Corpus Callosum |  | -8 | -42 | 18 | 6.88 | 2 |
| Suballosal Gyrus | 25 | -4 | 6 | -14 | 6.72 | 1 |
| Middle Frontal Gyrus |  | 26 | 34 | 42 | 6.61 | 1 |
| **FDR applied** |  |  |  |  |  |  |
| Posterior Cingulate Cortex | 5, 7, 24, 31 | -12 | -32 | 38 | 6.54 | 6905 |
| Middle Temporal Gyrus | 19, 22, 39, 40 | 66 | -38 | 24 | 5.75 | 1774 |
| Medial Frontal Gyrus | 9, 10, 32 | -4 | 52 | 10 | 6.54 | 965 |
| Superior Frontal Gyrus | 8, 9, 10 | 24 | 32 | 38 | 6.84 | 801 |
| Superior Frontal Gyrus | 8, 9, 10 | -24 | 44 | 26 | 5.42 | 743 |
| Inferior Parietal Lobule | 1, 2, 3, 40 | -62 | -28 | 30 | 7.38 | 637 |
| Culmen |  | -32 | -50 | -34 | 5.80 | 606 |
| Middle Temporal Gyrus | 19, 39 | -56 | -70 | 22 | 5.31 | 599 |
| Insula | 6, 13, 44 | -42 | 0 | 6 | 5.26 | 209 |
| Middle Frontal Gyrus | 6 | 22 | 2 | 64 | 3.99 | 95 |
| Anterior Cingulate Cortex | 11, 25 | -10 | 22 | -8 | 4.27 | 45 |
| Hippocampus |  | 20 | -40 | 10 | 4.43 | 44 |
| Pons |  | 18 | -34 | -36 | 4.10 | 37 |
| Thalamus |  | -12 | -26 | 6 | 4.02 | 37 |
| Parahippocampa Gyrus | 19, 36, 37 | 32 | -46 | -10 | 3.61 | 31 |
| Superior Frontal Gyrus | 10, 46 | -24 | 62 | 22 | 3.94 | 30 |
| Insula | 13 | 40 | 6 | -6 | 3.52 | 30 |
| Medial Frontal Gyrus | 11, 32 | 0 | 32 | -12 | 3.88 | 28 |
| Parahippocampa Gyrus |  | 44 | -28 | -18 | 3.56 | 27 |
| Superior Temporal Gyrus | 22 | 52 | 0 | 4 | 3.65 | 18 |
| Temporal Lobe | 20 | -40 | -14 | -24 | 4.15 | 17 |
| Insula | 13 | -38 | -22 | 30 | 3.72 | 16 |
| Caudate |  | 20 | 20 | 10 | 3.85 | 14 |
| Corpus Callosum |  | 2 | -6 | 18 | 3.71 | 12 |
| Parietal Lobe |  | 30 | -32 | 44 | 3.68 | 12 |
| Thalamus |  | 2 | -22 | 6 | 3.57 | 12 |
| Extra-Nuclear |  | 26 | -14 | 24 | 3.66 | 11 |
| Thalamus |  | 0 | -12 | 2 | 4.17 | 9 |
| Superior Frontal Gyrus | 10, 46 | -40 | 54 | 20 | 3.81 | 5 |
| Caudate |  | -14 | 20 | 10 | 3.69 | 3 |
| Temporal Lobe |  | -40 | -52 | 2 | 3.49 | 3 |
| Hippocampus |  | -18 | -40 | 6 | 3.37 | 2 |
| Superior Frontal Gyrus | 10 | 24 | 66 | 0 | 3.36 | 2 |
| Insula |  | 34 | -18 | 2 | 3.31 | 2 |
| Parahippocampal Gyrus |  | 28 | -42 | -4 | 3.37 | 1 |
| Extra-Nuclear |  | 24 | 12 | 14 | 3.35 | 1 |
| Caudate |  | 20 | 10 | 20 | 3.33 | 1 |
| Culmen |  | 0 | -50 | -4 | 3.32 | 1 |
| Postcentral Gyrus | 5 | -28 | -44 | 70 | 3.32 | 1 |
| Insula |  | 36 | 8 | 8 | 3.30 | 1 |
| Cluster-wise correction applied |  |  |  |  |  |  |
| Cingulate Gyrus | 5, 7, 24, 31 | -12 | -32 | 28 | 6.54 | 2297 |
| Posterior Cingulate Cortex | 7, 23, 30, 31 | 6 | -54 | 18 | 6.37 | 1205 |
| Medial Frontal Gyrus | 9, 10, 32 | -4 | 52 | 10 | 6.54 | 633 |
| Superior Frontal Gyrus | 8, 9, 10 | 24 | 32 | 38 | 6.84 | 495 |
| Middle Temporal Gyrus | 19, 23, 22, 39 | 46 | -72 | 18 | 5.63 | 459 |
| Inferior Parietal Lobule | 1, 2, 40 | -62 | -28 | 30 | 7.38 | 439 |
| Middle Frontal Gyrus | 8, 9, 10 | -24 | 44 | 26 | 5.42 | 437 |
| Inferior Parietal Lobule | 22, 40 | 66 | -38 | 24 | 5.75 | 432 |
| Middle Temporal Gyrus | 19, 39 | -56 | -70 | 22 | 5.31 | 397 |
| Culmen | 19, 37 | -32 | -50 | -34 | 5.80 | 340 |
| Anterior Cingulate Cortex | 24, 32 | 0 | 30 | 28 | 5.83 | 324 |
| SnPM |  |  |  |  |  |  |
| Inferior Parietal Lobule | 40 | -62 | -28 | 30 | 7.38 | 13 |
| Superior Frontal Gyrus |  | 24 | 32 | 38 | 6.84 | 6 |
| FDR applied |  |  |  |  |  |  |
| Medial Frontal Gyrus | 9, 10, 24, 32 | 8 | 52 | 16 | 8.34 | 2394 |
| Precuneus | 7, 23, 31 | -4 | -56 | 26 | 8.17 | 305 |
| Middle Temporal Gyrus | 20, 21 | 48 | 0 | -34 | 4.44 | 27 |
| Superior Frontal Gyrus | 8 | 26 | 40 | 48 | 5.10 | 23 |
| Medial Cingulate Cortex | 24 | 2 | -16 | 40 | 4.12 | 21 |
| Caudate |  | -16 | 22 | 8 | 4.81 | 16 |
| Angular Gyrus | 39 | -50 | -76 | 39 | 4.59 | 10 |
| Superior Temporal Gyrus | 39 | 58 | -64 | 22 | 4.09 | 6 |
| Extra-Nuclear |  | 14 | -4 | -10 | 4.03 | 4 |
| Superior Frontal Gyrus | 6 | 12 | 22 | 64 | 3.94 | 4 |
| Insula |  | -26 | 12 | -14 | 3.96 | 2 |
| Middle Frontal Gyrus |  | -22 | 34 | -14 | 3.93 | 2 |
| Medial Frontal Gyrus |  | -12 | 68 | 8 | 4.01 | 1 |
| Middle Temporal Gyrus | 9, 10, 24, 32 | -56 | -70 | 26 | 3.92 | 1 |
| **Cluster-wise correction applied** |  |  |  |  |  |  |
| Medial Frontal Gyrus | 9, 10, 32 | 8 | 52 | 16 | 8.34 | 2596 |
| Precuneus | 7, 23, 31 | -4 | -56 | 26 | 8.17 | 345 |
| **RFT-FWE applied** |  |  |  |  |  |  |
| Medial Frontal Gyrus | 9, 10 | 8 | 52 | 16 | 8.34 | 28 |
| Medial Frontal Gyrus | 10 | -2 | 62 | 2 | 8.06 | 9 |
| Posterior Cingulate Cortex | 31 | -4 | -56 | 26 | 8.17 | 8 |
| Medial Frontal Gyrus | 10 | -2 | 54 | -2 | 7.36 | 1 |
| **Bonferroni's FWE** |  |  |  |  |  |  |
| N/A |  |  |  |  |  |  |
| **SnPM** |  |  |  |  |  |  |
| Medial Frontal Gyrus | 9, 10, 32 | 10 | 50 | 16 | 9.66 | 141 |
| Medial Frontal Gyrus | 10 | -2 | 60 | 0 | 8.48 | 83 |
| Anterior Cingulate Cortex | 10, 32 | -10 | 40 | 4 | 8.23 | 77 |
| Anterior Cingulate Cortex | 24, 32 | -2 | 32 | 20 | 7.60 | 33 |
| Precuneus | 31 | -4 | -56 | 26 | 8.32 | 25 |
| Insula | 47 | -28 | 10 | -16 | 7.05 | 6 |
| Corpus Callosum |  | -10 | 20 | 18 | 7.16 | 3 |
| Middle Temporal Gyrus | 21 | 50 | 2 | -30 | 7.23 | 2 |
| **Moral Impersonal vs. Moral Personal** |  |  |  |  |  |  |
| **FDR applied** |  |  |  |  |  |  |
| Inferior Parietal Lobule | 7, 19, 39, 40 | 32 | -46 | 36 | 9.03 | 3301 |
| Inferior Parietal Lobule | 7, 19, 39, 40 | -28 | -60 | 46 | 6.55 | 2661 |
| Middle Frontal Gyrus | 6, 8, 9, 46 | -38 | 8 | 32 | 6.35 | 1706 |
| Declive |  | -4 | -76 | -28 | 7.01 | 1399 |
| Middle Temporal Gyrus | 20, 21, 22, 37 | -48 | -50 | -4 | 6.90 | 794 |
| Middle Temporal Gyrus | 20, 37 | 52 | 44 | -6 | 6.78 | 344 |
| Middle Frontal Gyrus | 6, 24, 32 | 24 | 6 | 54 | 4.87 | 289 |
| Extra-Nuclear |  | 24 | 12 | 34 | 4.78 | 226 |
| Inferior Frontal Gyrus | 8, 9 | 52 | 10 | 30 | 4.94 | 214 |
| Medial Frontal Gyrus | 6, 24, 32 | -12 | 8 | 52 | 4.52 | 123 |
| Insula | 13 | 30 | 24 | 13 | 4.52 | 66 |
| Middle Frontal Gyrus | 46 | 46 | 42 | 30 | 4.17 | 62 |
| Middle Frontal Gyrus | 10, 47 | -46 | 42 | -4 | 4.51 | 47 |
| Middle Occipital Gyrus | 18 | 28 | -84 | 0 | 3.66 | 45 |
| Middle Frontal Gyrus |  | 40 | 28 | 26 | 3.92 | 42 |
| Inferior Frontal Gyrus | 22, 44, 45 | -60 | 12 | 8 | 3.61 | 36 |
| Declive |  | -48 | -54 | -30 | 4.38 | 15 |
| Pons |  | 18 | -38 | -38 | 4.31 | 13 |
| Cuneus |  | -22 | -96 | 4 | 3.56 | 13 |
| Middle Frontal Gyrus | 10, 47 | 42 | 56 | 16 | 3.54 | 11 |
| Precentral Gyrus |  | -44 | 6 | 12 | 4.06 | 10 |
| Pons |  | 0 | -30 | -36 | 3.80 | 9 |
| Insula | 13 | -30 | 20 | 4 | 3.68 | 9 |
| Cingulate Gyrus | 32 | 6 | 20 | 46 | 3.59 | 6 |
| Middle Frontal Gyrus | 10, 47 | -38 | 58 | 14 | 3.66 | 5 |
| Temporal Lobe |  | 34 | -58 | -4 | 3.45 | 5 |
| Inferior Frontal Gyrus | 45, 47 | -58 | 32 | 0 | 3.60 | 4 |
| Superior Temporal Gyrus |  | -56 | -50 | 14 | 3.39 | 4 |
| Cuneus | 8 | 16 | -82 | 16 | 3.66 | 3 |
| Cuneus |  | 22 | -98 | -4 | 3.39 | 3 |
| Fusiform Gyrus | 20 | 36 | -38 | -20 | 3.38 | 3 |
| Temporal Lobe |  | 30 | -44 | 0 | 3.66 | 2 |
| Orbitofrontal Cortex |  | 46 | 46 | -16 | 3.66 | 1 |
| Middle Frontal Gyrus | 10, 47 | -30 | 64 | 14 | 3.51 | 1 |
| Middle Frontal Gyrus | 10, 47 | -32 | 62 | 16 | 3.42 | 1 |
| Extra-Nuclear |  | 28 | 28 | 4 | 3.34 | 1 |
| **Cluster-wise correction applied** |  |  |  |  |  |  |
| Inferior Parietal Lobule | 7, 19, 39, 40 | 32 | -46 | 36 | 9.03 | 2621 |
| Inferior Parietal Lobule | 7, 19, 40 | -28 | -60 | 46 | 6.55 | 1979 |
| Inferior Frontal Gyrus | 6, 8, 9, 46 | -38 | 8 | 32 | 6.35 | 991 |
| Declive |  | -4 | -76 | -28 | 7.01 | 768 |
| Middle Temporal Gyrus | 19, 21, 22, 37 | -48 | -50 | -4 | 6.90 | 467 |
| **RFT-FWE applied** |  |  |  |  |  |  |
| Angular Gyrus |  | 32 | -46 | 36 | 9.03 | 22 |
| **Bonferroni's FWE** |  |  |  |  |  |  |
| Parietal Lobe |  | 32 | -46 | 36 | 9.03 | 2 |
| **SnPM** |  |  |  |  |  |  |
| Angular Gyrus | 7, 19, 39, 40 | 32 | -46 | 36 | 9.03 | 203 |
| Declive |  | -4 | -76 | -28 | 7.01 | 10 |
| Temporal Lobe |  | -48 | -50 | -4 | 6.90 | 6 |
| Temporal Lobe |  | 52 | -44 | -6 | 6.78 | 6 |
| Fastigium |  | 8 | -58 | -26 | 6.70 | 5 |
| Middle Temporal Gyrus | 20 | 56 | -48 | -14 | 6.56 | 3 |
| Superior Parietal Lobule |  | -28 | -60 | 46 | 6.55 | 2 |

**Figure S1.**
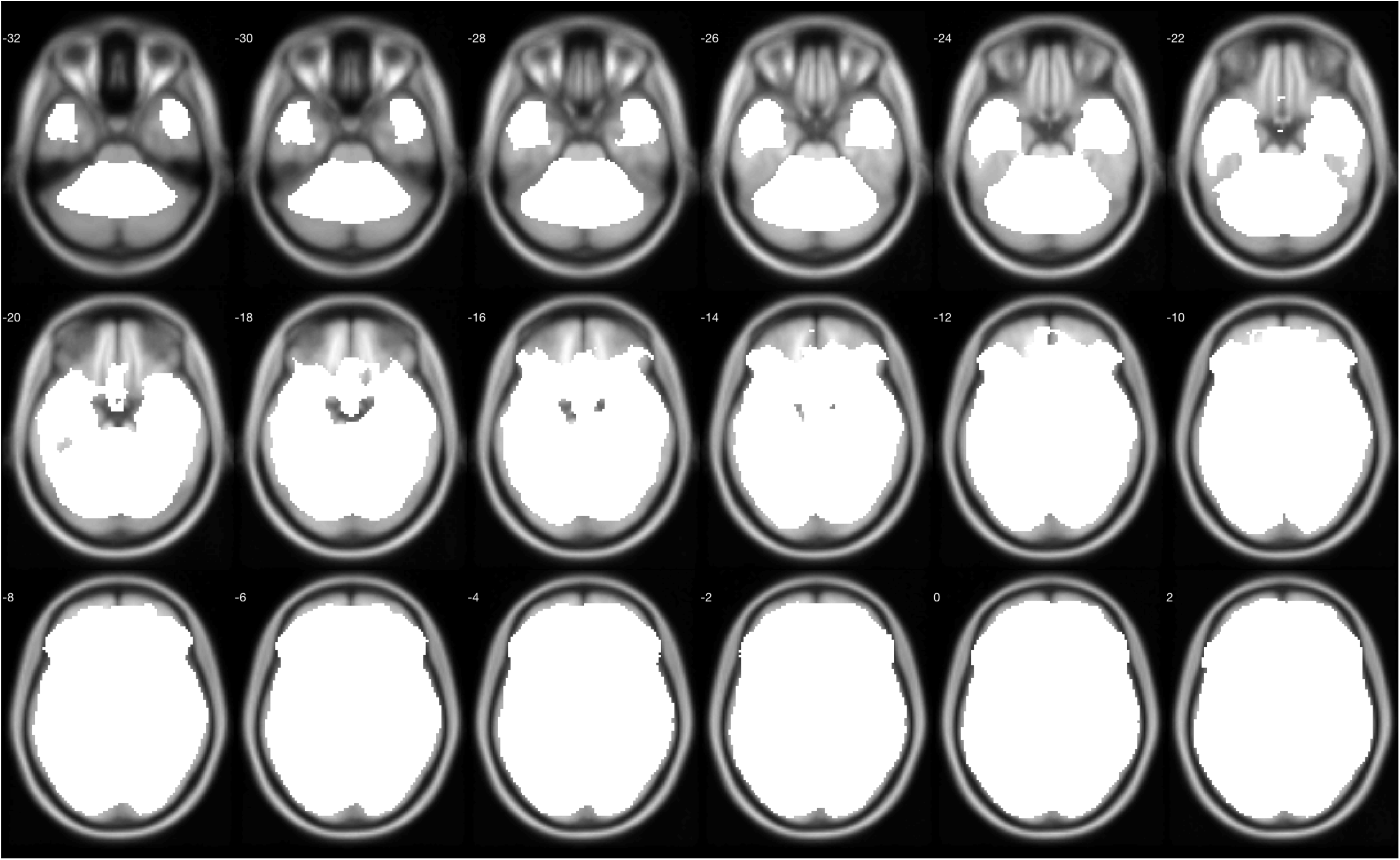
The implicit mask in SPM near the VMPFC.

